# Ventral temporal and posteromedial sulcal morphology in autism spectrum disorder

**DOI:** 10.1101/2022.09.01.506213

**Authors:** Javier Ramos Benitez, Sandhya Kannan, William L. Hastings, Benjamin J. Parker, Ethan H. Willbrand, Kevin S. Weiner

**Author notes:** Corresponding author:* Kevin S. Weiner. equal contribution (Javier Ramos Benitez, Sandhya Kannan). Author contributions:* J.R.B., S.K., E.H.W., K.S.W. designed research; J.R.B., S.K., W.L.H., B.J.P., E.H.W., and K.S.W. performed manual sulcal labeling; J.R.B., S.K., E.H.W., K.S.W. analyzed data; J.R.B., S.K., E.H.W., K.S.W. wrote the paper; all authors edited the paper and gave final approval before submission.

## Abstract

Two recent parallel research tracks link tertiary sulcal morphology—sulci that emerge last in gestation and continue to develop after birth—with functional features of the cerebral cortex and cognition, respectively. The first track identified a relationship between the mid-fusiform sulcus (MFS) in ventral temporal cortex (VTC) and cognition in individuals with Autism Spectrum Disorder (ASD). The second track identified a new tertiary sulcus, the inframarginal sulcus (IFRMS), that serves as a tripartite landmark within the posteromedial cortex (PMC). As VTC and PMC are structurally and functionally different in individuals with ASD compared to neurotypical controls (NTs), here, we integrated these two tracks with a twofold approach. First, we tested if there are morphological differences in VTC and PMC sulci between 50 NTs and 50 individuals with ASD. Second, we tested if tertiary sulcal morphology was linked to cognition in ASD individuals. Our twofold approach replicates and extends recent findings in five ways. First, in terms of replication, the standard deviation (STD) of MFS cortical thickness (CT) was increased in ASDs compared to NTs. Second, MFS length was shorter in ASDs compared to NTs. Third, the CT STD effect extended to other VTC and PMC sulci. Fourth, a subset of VTC and PMC morphological features were correlated between regions in ASD. Fifth, IFRMS depth was negatively associated with ADOS-GS score. These results empirically support a relationship between later-developing, tertiary sulci and ASD, providing a novel framework to study the relationship between brain structure and cognition in additional neurodevelopmental disorders in future studies.

**Lay Summary:** We observed that some, but not all, morphological features of later-developing tertiary indentations (sulci) in the cerebral cortex differed significantly between neurotypical controls and individuals with autism spectrum disorder (ASD). In ASD, a subset of sulcal morphological features also correlated between brain areas and one feature reflected differences in cognition. Thus, studying these structures provides insight into how individual variability in structure is related to individual variability in cognition in ASD.

## Introduction

Understanding neuroanatomical features contributing to neurodevelopmental disorders is a major goal in cognitive and clinical neuroscience. Previous studies have identified cortical morphological differences between individuals with Autism Spectrum Disorder (ASD) and neurotypical controls (NTs), such as cortical thickness (Bethlehem et al., 2020; Hardan, Muddasani, Vemulapalli, Keshavan, & Minshew, 2006; Khundrakpam, Lewis, Kostopoulos, Carbonell, & Evans, 2017; Nunes et al., 2020; Smith et al., 2016; Yang, Beam, Pelphrey, Abdullahi, & Jou, 2016; Zielinski et al., 2014) and gray matter volume (Guo et al., 2021; Hazlett, Poe, Gerig, Smith, & Piven, 2006; Prigge et al., 2021; Sato et al., 2017; Seng, Lai, Goh, Tseng, & Gau, 2022; Yamasaki et al., 2010). Yet, very few have considered tertiary sulci, which emerge last in gestation and show a protracted development after birth (Armstrong, Schleicher, Omran, Curtis, & Zilles, 1995; Chi, Dooling, & Gilles, 1977; Petrides, 2019; Weiner, 2019; Welker, 1990). Most relevant to the present study, two recent studies have implicated a role of tertiary sulci in neurodevelopmental disorders in two ways. First, tertiary sulci in high-level visual cortex were shorter in individuals with developmental prosopagnosia (DP) compared to neurotypical controls (NTs), and individual differences in MFS length also predicted individual differences in face perception ability (Parker et al., 2022). Second, the cortical thickness of tertiary sulci in high-level visual cortex in ASD individuals was recently shown to be negatively correlated with a behavioral task that tests an individual’s ability to interpret mental states and emotions from facial features (Ammons et al., 2021). Given these previous findings, the present study first aimed to replicate and extend these findings by comparing the morphology of VTC sulci between ASD and NT participants (as in Ammons et al., 2021). Here, we also considered additional VTC sulci and morphological features not previously studied.

In addition to VTC, previous work shows that the posteromedial cortex (PMC) is structurally and functionally different in individuals with ASD compared to NTs (Haghighat, Mirzarezaee, Araabi, & Khadem, 2021; Leech & Sharp, 2014; Lynch et al., 2013; Oblak, Gibbs, & Blatt, 2010; Oblak, Rosene, Kemper, Bauman, & Blatt, 2011). Additionally, previous studies found cytoarchitectural differences in the PMC of individuals with ASD compared to NTs (Oblak et al., 2011), as well as decreased binding sites for GABAergic receptors in the superficial and deep layers of PMC in ASD individuals compared to NTs (Oblak et al., 2010). Crucially, recent work has clarified the sulcal organization of PMC and uncovered the presence of a novel, consistent tertiary sulcus—the inframarginal sulcus (IFRMS)—that is related to the functional and anatomical organization of PMC (Willbrand et al., in press). Accordingly, these PMC sulci have yet to be characterized in ASD individuals. To fill this gap in knowledge, we compared the morphological features of PMC sulci between ASD and NT participants.

Together, the goals of the present study were fivefold: i) attempt to replicate previous work (Ammons et al., 2021) and to quantitatively compare VTC sulcal morphology between individuals with ASD and NTs; ii) to extend these analyses to other VTC sulci and morphological features not previously studied; iii) to assess whether these morphological similarities and differences extend to PMC sulci; iv) to determine if the morphology of VTC and PMC tertiary sulci are correlated in ASD and/or NT participants; and v) to determine if individual differences in the morphology of VTC and/or PMC tertiary sulci are related to individual differences in Autism Diagnosis Observation Schedule-General Social (ADOS-GS) scores. To our knowledge, this is the first study to examine (i) if tertiary sulci in multiple cortical expanses differ between individuals with ASD and NTs and (ii) if individual differences in PMC tertiary sulcal morphology relate to individual differences in cognition.

## Materials and Methods

This study utilized data from the Autism Brain Imaging Data Exchange (ABIDE; (Di Martino et al., 2014). Below, we detail each aspect of this multimodal dataset used for the present study.

### Participants

Here we used data from 100 randomly chosen individuals in the ABIDE dataset (http://fcon_1000.projects.nitrc.org/indi/abide/abide_II.html). Data were from New York University (N = 78) and Georgetown University (N = 22) collection sites. One half of the data (N = 50) were from individuals with Autism Spectrum Disorder (ASD) and the other half of the data (N = 50) were from neurotypical controls (NTs). The age range for ASDs (ages 5 to 34) and NTs (ages 5 to 24) was comparable (*p* = .87). See Supplementary Table 1 for additional demographic details. Inclusion criteria for ASDs was a clinician’s diagnosis based on the DSM-IV or DSM-V.

### Neuroanatomical data

#### Imaging acquisition

All children who participated in the study completed a mock scanning session in order to become acclimated to the scanning environment. Standard T1-weighted MPRAGE anatomical scans (TR = 2530 ms, TE = 3.25 ms, 1.3 × 1 × 1.3 mm^3^ voxels) were acquired for cortical morphometric analyses in these participants using a 3T Siemens Allegra MRI scanner at New York University Langone Medical Center and the Center for Functional and Molecular Imaging at Georgetown University.

#### Cortical surface reconstruction

Each T1-weighted image was inspected for scanner artifacts. Cortical surface reconstructions were generated for each participant from the aforementioned scans using a standard FreeSurfer pipeline (Dale, Fischl, & Sereno, 1999; Fischl, Sereno, & Dale, 1999; Fischl, Sereno, Tootell, & Dale, 1999); FreeSurfer (v6.0): surfer.nmr.mgh.harvard.edu/). Cortical surface reconstructions were examined for segmentation errors and manually corrected when necessary. Sulcal labels and the anatomical metrics extracted from those labels were calculated on each cortical surface in each participant’s native space.

### Regions of interest (ROIs) and sulcal labeling procedure

#### Ventral Temporal Cortex

As in our previous work across ages (Weiner et al., 2014), species (Miller et al., 2020), and individuals with Developmental Prosopagnosia [2% of the population who cannot perceive faces, but do not have brain damage; (Parker et al., 2022)], three sulci were manually identified in the VTC for each hemisphere: the Collateral Sulcus (CoS), Occipito-Temporal Sulcus (OTS), and Mid-Fusiform Sulcus (MFS). A three-tiered approach was implemented to manually label each of these three sulci in every hemisphere. First, the FG was identified as the major gyrus in VTC. Second, once the FG was identified, the OTS, CoS, and MFS were identified based on the following criteria: 1) the CoS was identified as a deep and long sulcus identifying the medial extent of the FG, 2) the OTS was identified as a deep and long sulcus identifying the lateral extent of the FG, and 3) the MFS was identified as either a single shallow longitudinal sulcus dividing the FG into lateral and medial partitions or it was identified as two or more shallow sulcal components dividing the FG into lateral and medial partitions. Third, as the MFS has predictable anterior (posterior extent of the hippocampus) and posterior (posterior transverse collateral sulcus) landmarks, but the OTS and CoS can extend longitudinally from the occipital pole to the temporal pole, we restricted our OTS and CoS definitions to the portions in VTC surrounding the MFS. This is consistent with our previous protocols and assures that the portions of the OTS and CoS being compared to the MFS are within the VTC and are approximately equal along the length of the three sulci. Further, this restriction assures that any morphological differences between groups are not due to the fact that the OTS and CoS are much longer than the MFS, for example. Example hemispheres are shown in **Figure 1** (Supplementary Figs. 1 and 2 for all participants).

**Figure 1.**
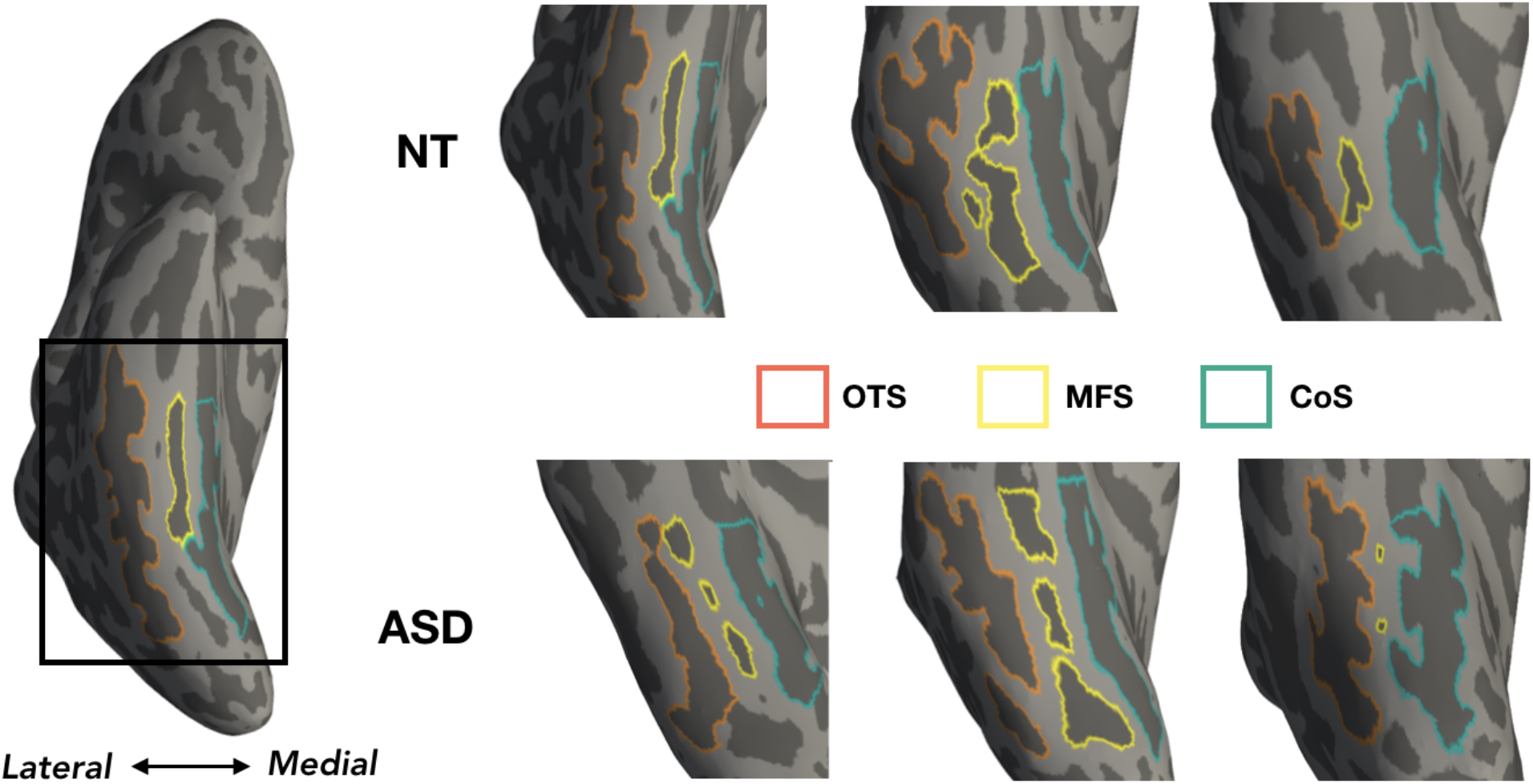
Sulci of interest in ventral temporal cortex (VTC). *Left:* One example inflated cortical surface indicating the location (rectangle) of the zoomed portions indicated on the right. *Right:* Three example inflated cortical surface reconstructions of right hemispheres from NTs (top) and individuals with ASD (bottom). The leftmost hemisphere displays the mid-fusiform sulcus (MFS) at mean length, the center at max length, and the rightmost at minimum length in each respective group. The MFS is outlined in yellow, the collateral sulcus (CoS) in teal, and the occipito-temporal sulcus (OTS) in red. Dark gray: sulci. Light gray: gyri.

#### Posteromedial Cortex (PMC)

Within the PMC, we identify sulci based on the recent proposal by Petrides (Petrides, 2019) and recent work in individual participants that builds on this work (Willbrand et al., in press). The PMC consists of the precuneus (PrC), posterior cingulate cortex (PCC), and retrosplenial cortex [RSC; (Foster & Parvizi, 2017; Parvizi, Van Hoesen, Buckwalter, & Damasio, 2006; Willbrand et al., in press)]. Based on prior work (Willbrand et al., in press), the PrC can be defined by the following anterior, posterior, and inferior boundaries in the medial parietal cortex: the marginal ramus of the cingulate sulcus (MCGS), the parieto-occipital sulcus (POS), and splenial sulcus (SPLS), respectively. Conversely, the PCC is bounded by the cingulate sulcus (CGS), MCGS, and SPLS superiorly, callosal sulcus (CAS) inferiorly, and POS posteriorly.

Within the PRC, we labeled the precuneual limiting sulcus (PRCULS, which extends from the superior portion of the POS), the precuneal sulci (PRCUS), and the superior parietal sulcus (SPS). Specifically, while previous studies identify one to three PRCUS (Margulies et al., 2009; Ono, Kubik, & Abernathey, 1990; Petrides, 2019; Vogt, Nimchinsky, Vogt, & Hof, 1995; Vogt, Vogt, & Laureys, 2006), we recently identified three precuneal sulci in every hemisphere (Willbrand et al., in press). The PRCUS were labeled based on their position within the PrC: posterior precuneal (PRCUS-p), intermediate precuneal (PRCUS-i), and anterior precuneal (PRCUS-a) sulci. As in prior work (Willbrand et al., in press), we also identified the SPS when it was present entering the PrC. Within the PCC, we also defined three tertiary sulci that were recently identified. First, the inframarginal sulcus (IFRMS) is a newly identified sulcus shown to be a tripartite landmark in the human brain (Willbrand et al., in press). The IFRMS is typically present in every hemisphere inferior to the MCGS and anterior to the SPLS. In addition, we identify two other tertiary sulci (which are more variable across hemispheres and not identifiable in every single hemisphere): the subsplenial sulcus (SSPLS) which is inferior to the SPLS and the posterior intracingulate sulcus (ICGS-p) which is anterior to the IFRMS. Example hemispheres are shown in **Figure 2** (Supplementary Figs. 3 and 4 for all participants).

**Figure 2.**
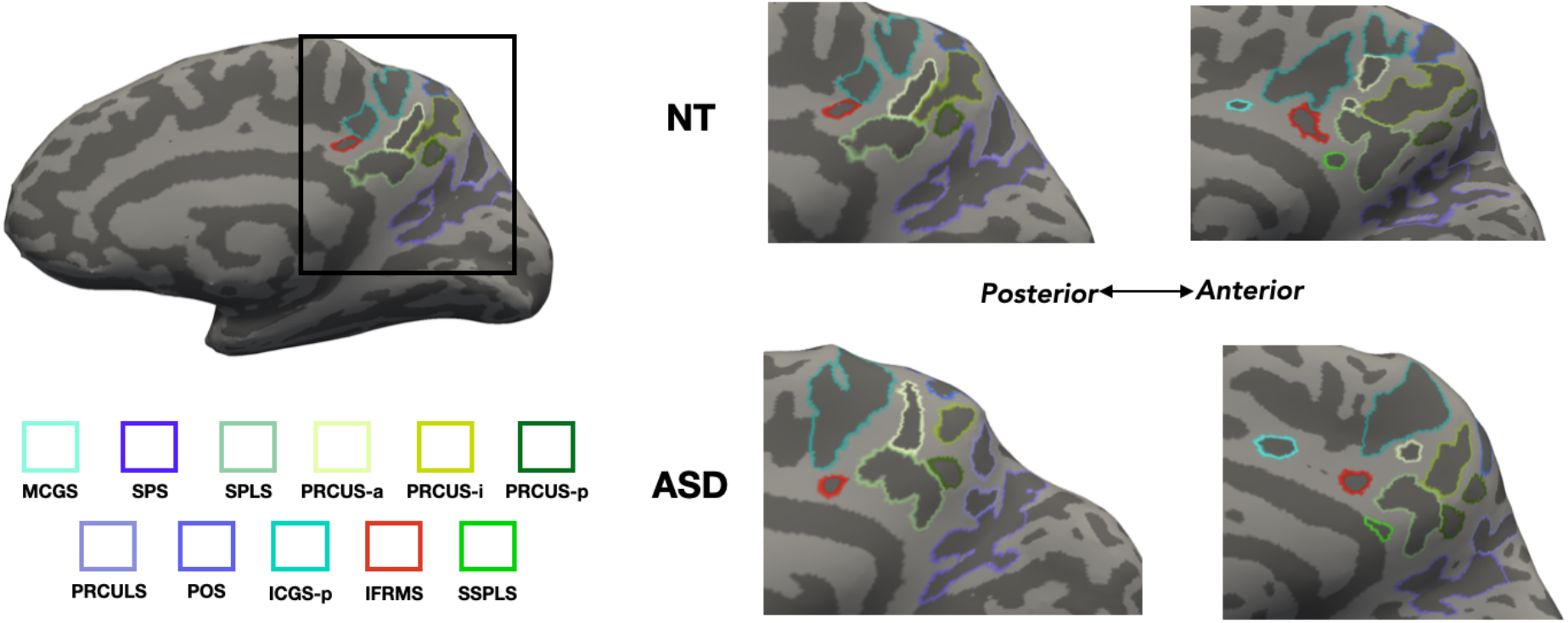
Sulci of interest in posteromedial cortex (PMC). *Left:* One example inflated cortical surface indicating the location (rectangle) of the zoomed portions indicated on the right. *Right:* Two example inflated cortical surface reconstructions of right hemispheres from NTs (top) and individuals with ASD (bottom). The leftmost images display right hemispheres with only the ifrms present, and the rightmost images display the variable sulci present in the PMC in both groups (sspls, green; icgs-p, teal).

#### Manual sulcal labeling

Sulci were manually defined for each individual hemisphere, blind to each group, with tksurfer tools on the FreeSurfer inflated view as described in previous publications (Miller, Voorhies, Lurie, D’Esposito, & Weiner, 2021; Voorhies, Miller, Yao, Bunge, & Weiner, 2022). *Pial, inflated*, and *smooth* surfaces were used to determine the boundaries between intersecting sulci. Sulcal labels were defined in a two-step system as in our previous work (Miller et al., 2021; Voorhies et al., 2021). First, sulcal labels were defined manually by trained raters (J.R., S.K., W.H., B.J.P., E.H.W.) and then revised and finalized by a neuroanatomist (K.S.W.). The label definition process was completed in a blind fashion in all hemispheres before conducting morphological analyses of sulcal labels within and between groups.

### Morphological Analyses

#### Extraction of morphological features

Considering that shallow depth and smaller surface area (SA) are defining characteristic features of tertiary sulci compared to primary and secondary sulci, our main focus was on mean sulcal depth and total SA. We also calculated the max sulcal depth (known as the sulcal pit), which is considered the first point of a sulcus to pull in development and considered to be particularly relevant in the development of functional regions in gyrencephalic brains (Natu et al., 2021; Rakic, 1988; Régis et al., 2005). Additionally, we also considered mean cortical thickness (CT; mm) and total gray matter volume (GMV; mm^3^), which are common morphological features explored in previous neuroimaging studies. Finally, we also quantified the standard deviation of cortical thickness (CT STD; mm) in each group as a recent study showed that the CT STD of the MFS was greater in ASDs compared to NTs (Ammons et al., 2021) with a comparable sample size. Mean and max sulcal depth values (in standard FreeSurfer units) were computed in native space from the .sulc file generated in FreeSurfer (Dale, Fischl, & Sereno, 1999). Briefly, depth values are calculated based on how far removed a vertex is from what is referred to as a “mid-surface,” which is determined computationally so that the mean of the displacements around this “mid-surface” is zero. Thus, generally, gyri have negative values, while sulci have positive values. Given the shallowness and variability in the depth of tertiary sulci (Voorhies et al., 2021; Yao, Voorhies, Miller, Bunge, & Weiner, 2022), some mean depth values extend below zero. We emphasize that this just reflects the metric implemented in FreeSurfer. All other morphological characteristics were calculated using the *mris_anatomical_stats* function (Fischl & Dale, 2000).

Given recent findings showing that a) the MFS is shorter in individuals with developmental prosopagnosia compared to NTs and b) MFS length is correlated with face perception ability (Parker et al., 2022), we also calculated MFS length (mm) for each participant. As in our previous work (Miller et al., 2020; Parker et al., 2022), we calculated length as the longest geodesic distance along the cortical surface between any pair of vertices on the border of the MFS label. When sulci were identified as consisting of multiple disconnected pieces on the cortical surface, the sulcal length was defined as the total length of each sulcal component [excluding the annectant gyral component(s)]. Here, geodesic distance was calculated on the fiducial surface using algorithms implemented in the Pycortex software package (https://gallantlab.github.io/).

#### Morphological comparisons between ASD and NT participants

All statistical analyses were performed using R (v4.0.4) and RStudio (v1.4). In addition to basic R functions, the additional packages *stats, car, rstatix, ggplot2, dplyr, emmeans*, and *sjstats* were utilized to make use of dedicated functions for graphing, data processing, and analysis. We used ANCOVAs to investigate differences between groups while also controlling for age (see Supplementary Table 1 for age range). ANCOVAs were implemented using functions from the built-in R *stats* package and the *car* package.

We investigated whether VTC sulcal morphology differed between groups with ANCOVAs using group (ASD, NT), sulcus (MFS, CoS, OTS), and hemisphere (left, right) as factors for sulcal depth (mean, max), GMV, SA, and CT (mean, STD). Age was included as a covariate. This analysis was primarily run to replicate and extend findings from Ammons et al. (2021). Additionally, to further extend this recent work (Ammons et al., 2021), we investigated group differences in MFS length using a 2-way ANCOVA (controlling for age) with group and hemisphere as factors.

Considering the large number of sulci in PMC compared to VTC (11 versus 3, respectively), we next assessed for differences between groups and sulcal types (non-tertiary vs. tertiary) in PMC using ANCOVAs (controlling for age) with group, sulcal type, and hemisphere as factors for sulcal depth (mean, max), GMV, SA, and CT (mean, STD).

We also performed an exploratory analysis to test if morphological features of VTC tertiary sulci (MFS) were correlated with PMC tertiary sulci (IFRMS, SSPLS, ICGS-p) with the hypothesis that since tertiary sulci emerge last in gestation, morphological features of tertiary sulci in different cortical expanses may be correlated. In all analyses, *p* < 0.05 was considered significant after controlling for multiple comparisons with the false discovery rate (FDR) method.

### Behavioral Analyses

Here, we focused on the Autism Diagnostic Observation Schedule (ADOS), which is a common diagnostic tool consisting of four modules designed to adjust for the language level and age of the individual tested to better capture the severity of symptoms across cognitive levels (Lord et al., 2006). Each subtest has different thresholds that may support an ASD diagnosis, and scores in each module vary across ages. For the purpose of analyzing behaviors related to facial processing, we analyzed the ADOS-General Social (ADOS-GS). This subtest contains items examining aspects of facial processing such as directed facial expressions and unusual eye contact, as well as social correlates that depend on visual processing such as social overtures and social responses. For this subtest, the average scores among those individuals in the autism spectrum are expected to be between 6-8 depending on age while some individuals may achieve the maximum of 14 points (ASD participants with ADOS-GS scores = 40; Mean ± STD = 7 ± 2.30).

Correlations were calculated between the morphology of the most consistent tertiary sulci (MFS and IFRMS) in each hemisphere and ADOS-GS scores in participants with ASD. Specifically, length (MFS), CT (MFS, IFRMS), and sulcal depth (MFS, IFRMS) were assessed given recent findings between these morphological features and cognition in NTs and patient populations for the MFS and other tertiary sulci [CT: (Ammons et al., 2021); SD: (Brun et al., 2016; Voorhies et al., 2021); length: (Garrison et al., 2015; Parker et al., 2022)].

## Results

### Greater variability in the CT of VTC sulci in ASD individuals compared to NTs

As in prior studies (Miller et al., 2020; Parker et al., 2022; Weiner et al., 2014), we could identify the MFS, CoS, and OTS in all participants in both hemispheres in both groups (**Fig. 1**; Supplementary Table 2; see Supplementary Figs. 1 and 2 for all 600 defined VTC sulci across 100 participants). For each morphological feature of interest [sulcal depth (mean, max), gray matter volume (GMV), surface area (SA), and cortical thickness (CT; mean, STD)], we ran a 3-way ANCOVA (controlling for age) with group (ASD, NT), sulcus (MFS, OTS, CoS), and hemisphere (left, right) as factors. Replicating Ammons and colleagues (2021), there was a main effect of group (F(1,587) = 8.70, *p* = .018, η^2^G = 0.015) for CT STD such that there was greater variance in the CT of VTC sulci across hemispheres in ASD individuals compared to NTs (**Fig. 3**). There were no other statistically significant group differences (main effects or interactions) for the other morphological features of interest (mean sulcal depth, max sulcal depth, SA, GMV, and mean CT) for VTC sulci (*p*s > .59; Supplementary Fig. 5).

**Figure 3.**
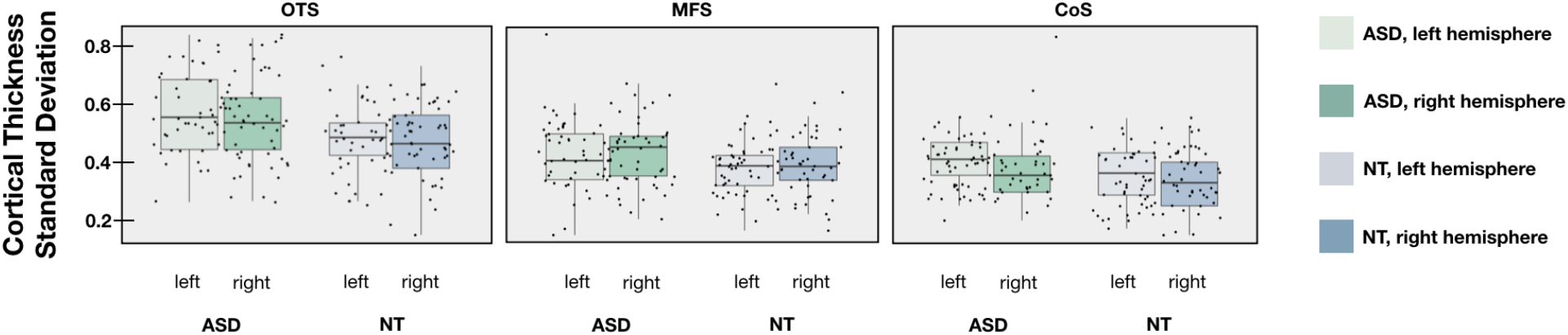
The thickness of VTC sulci is more variable in individuals with ASD compared to NTs. Boxplots indicating the standard deviation (± quartile) of sulcal cortical thickness in each group (ASD; NT) for the OTS (left), MFS (middle), and CoS (right). *Left:* ASD. *Right:* NT.

### The MFS is shorter in ASD individuals compared to NTs

Given recent work identifying a shorter MFS length in individuals with developmental prosopagnosia compared to NTs (Parker et al., 2022), we tested whether the length of the MFS (in mm) differed between ASD and NT participants using a 2-way ANCOVA (controlling for age) with group (ASD, NT) and hemisphere (left, right) as factors. There was a main effect of group (F(1,195) = 6.86, *p* = .03, η^2^G = 0.034), with no hemisphere interaction (*p* = .71), in which individuals with ASD had a shorter MFS than NTs in both hemispheres (**Fig. 4**; ASD Mean ± STD = 31 ± 10.5 mm; NT Mean ± STD = 38 ± 11.7 mm; Mean Difference = 7 mm).

**Figure 4.**
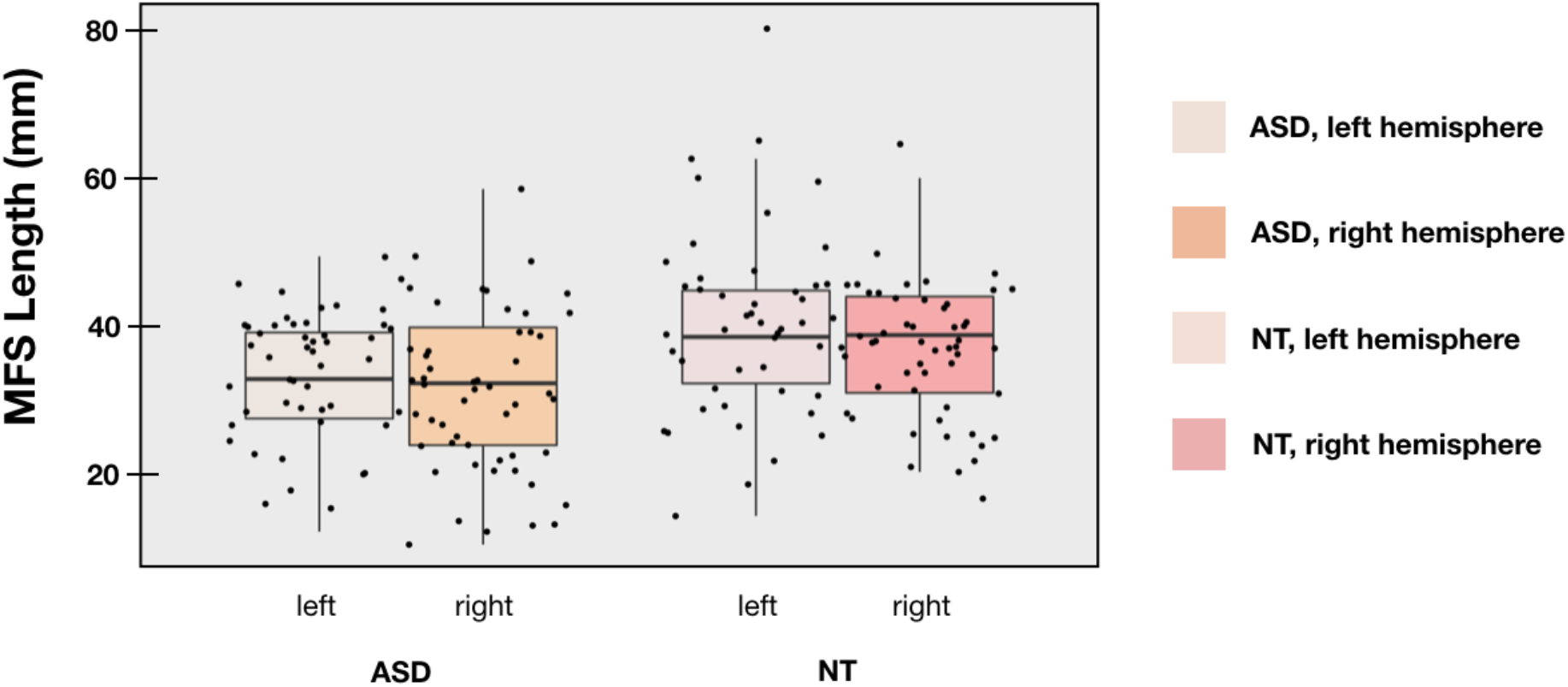
The MFS is shorter in ASD individuals compared to NTs. Boxplots indicating MFS length (± quartile) in each group. The left two plots indicate the left and right hemispheres of individuals with ASD and the right two plots indicate the left and right hemispheres of NTs.

### Greater variance in the CT of PMC sulci in ASD individuals compared to NTs

Similar to findings by Willbrand et al. (in press), the three sulci bounding the PMC—the MCGS, POS, and SPLS—were identifiable in all hemispheres and groups in our sample (Supplementary Table 2). The majority of PMC sulci were also present within all hemispheres and groups (Supplementary Table 2). In addition, the SPS was present in the majority of hemispheres in both groups (Supplementary Table 2). Within PCC, the IFRMS was identifiable in all but one left and right hemisphere, whereas the other two PCC sulci (SSPLS and ICGS-p) were more variable (Supplementary Table 2; see Willbrand et al., in press; Supplementary Figs. 3 and 4 for all 1979 PMC sulci defined across the 100 participants).

Comparable to VTC, 3-way ANCOVAs (controlling for age) with group (ASD, NT), sulcal type (non-tertiary, tertiary), and hemisphere (left, right) as factors revealed a significant main effect of group for CT STD (F(1,1970) = 19.95, *p* < .001, η^2^G=0.01) such that for both PMC sulcal types, the variance in CT was greater in individuals with ASD compared to NTs across hemispheres (**Fig. 5**). There were no other significant main effects or interactions for mean sulcal depth, max sulcal depth, GMV, SA, or mean CT (*p*s > .47; Supplementary Figs. 6-10).

**Figure 5.**
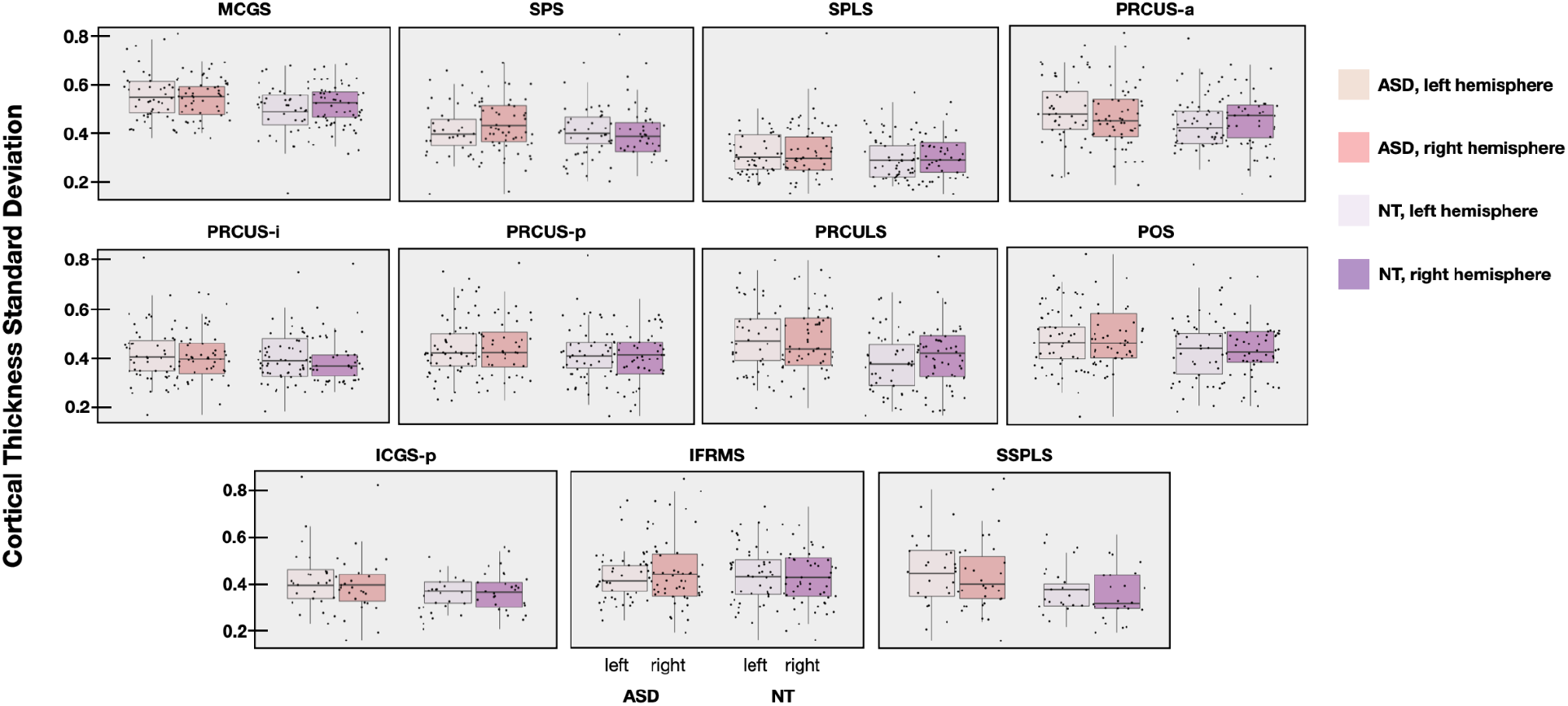
The thickness of PMC sulci is more variable in individuals with ASD compared to NTs. Boxplots indicating the standard deviation (± quartile) of sulcal cortical thickness in each group (ASD; NT) for the 11 PMC sulci. *Left:* ASD. *Right:* NT.

### Some, but not all, tertiary sulcal morphological features of VTC and PMC are correlated in ASD individuals, but not NTs

Recent findings proposed that tertiary sulci could be novel targets for neuropsychiatric disorders since they are largely specific to the hominoid brain and are related to human-specific aspects of cognition (Miller, D’Esposito, & Weiner, 2021; Parker et al., 2022). Thus, we tested if the morphology of the MFS in VTC was correlated with morphological features of PMC tertiary sulci (IFRMS, SSPLS, ICGS-p). This analysis revealed that some, but not all, tertiary sulcal morphological features of VTC and PMC are correlated in ASD individuals. For example, in the right hemisphere of ASD individuals, the mean sulcal depth of the MFS and IFRMS were significantly negatively correlated (*r* = - 0.37, *p* = 0.008; **Fig. 6A**), while in the left hemisphere of ASD individuals who had the SSPLS, the mean CT of the MFS and the SSPLS were correlated (*r* = 0.45, *p* = 0.01; **Fig. 6B**).

**Figure 6.**
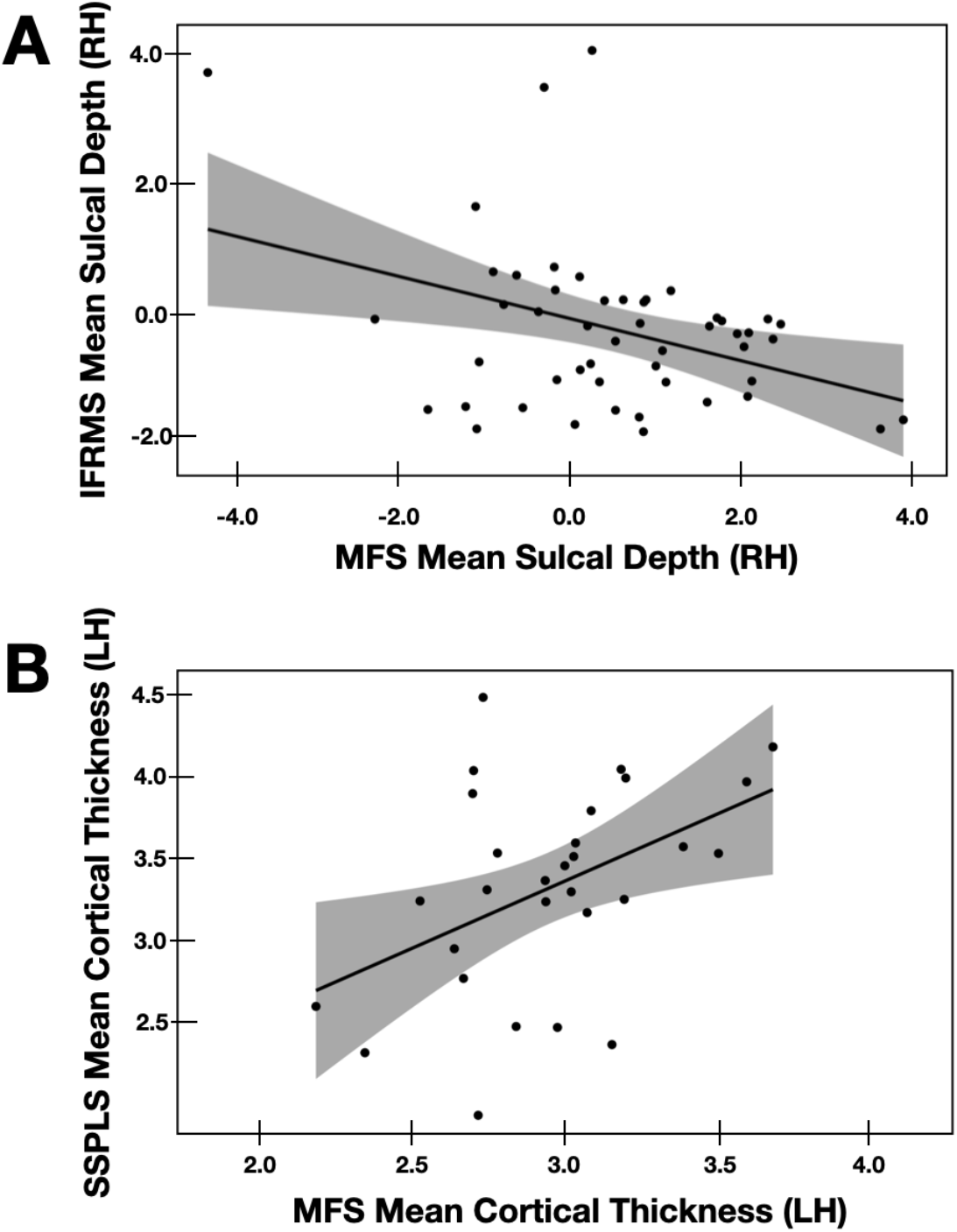
Some, but not all, tertiary sulcal morphological features of VTC and PMC are correlated in individuals with ASD, but not NTs. **A**. The mean sulcal depth of the MFS is negatively correlated with the mean sulcal depth of the IFRMS in the right hemisphere (*r* = -0.37, *p* = 0.0081). **B**. The mean cortical thickness of the MFS is positively correlated with the mean cortical thickness of the SSPLS in the left hemisphere (*r* = 0.45, *p* = 0.015).

### The morphology of the ifrms is related to cognition in ASD

Given the recent findings relating the morphology of later-developing tertiary sulci to later-developing, higher-level cognitive abilities, we tested if the morphology of the most consistent tertiary sulci in VTC and PCC (i.e., the MFS and IFRMS) predicted scores on the General Social of the Autism Diagnostic Observation Schedule [ADOS-GS; (Lord et al., 2006)], a subtest that measures eye contact and facial orientation (Methods). To do so, we included participants with ASD who also had ADOS-GS scores (N = 40). In these analyses, we focused on mean and max sulcal depth, mean and STD CT, and length (for the MFS), due to recent findings between these morphological features and cognition, specifically the sulcal depth of tertiary sulci in lateral PFC to reasoning (Voorhies et al., 2021), the CT (mean and STD) of the MFS to theory of mind in ASD (Ammons et al., 2021), and the length of the MFS to face perception ability (Parker et al., 2022). We found that the sulcal depth of the IFRMS in the left hemisphere was negatively related to ADOS-GS scores for both the mean (*r* = -.33, *p* = 0.035; **Fig. 7A**) and max (r = -.34, *p* = 0.031; **Fig. 7B**) sulcal depth. No other morphological feature of the IFRMS related to ADOS-GS scores in either hemisphere (*p*s > .22).

**Figure 7.**
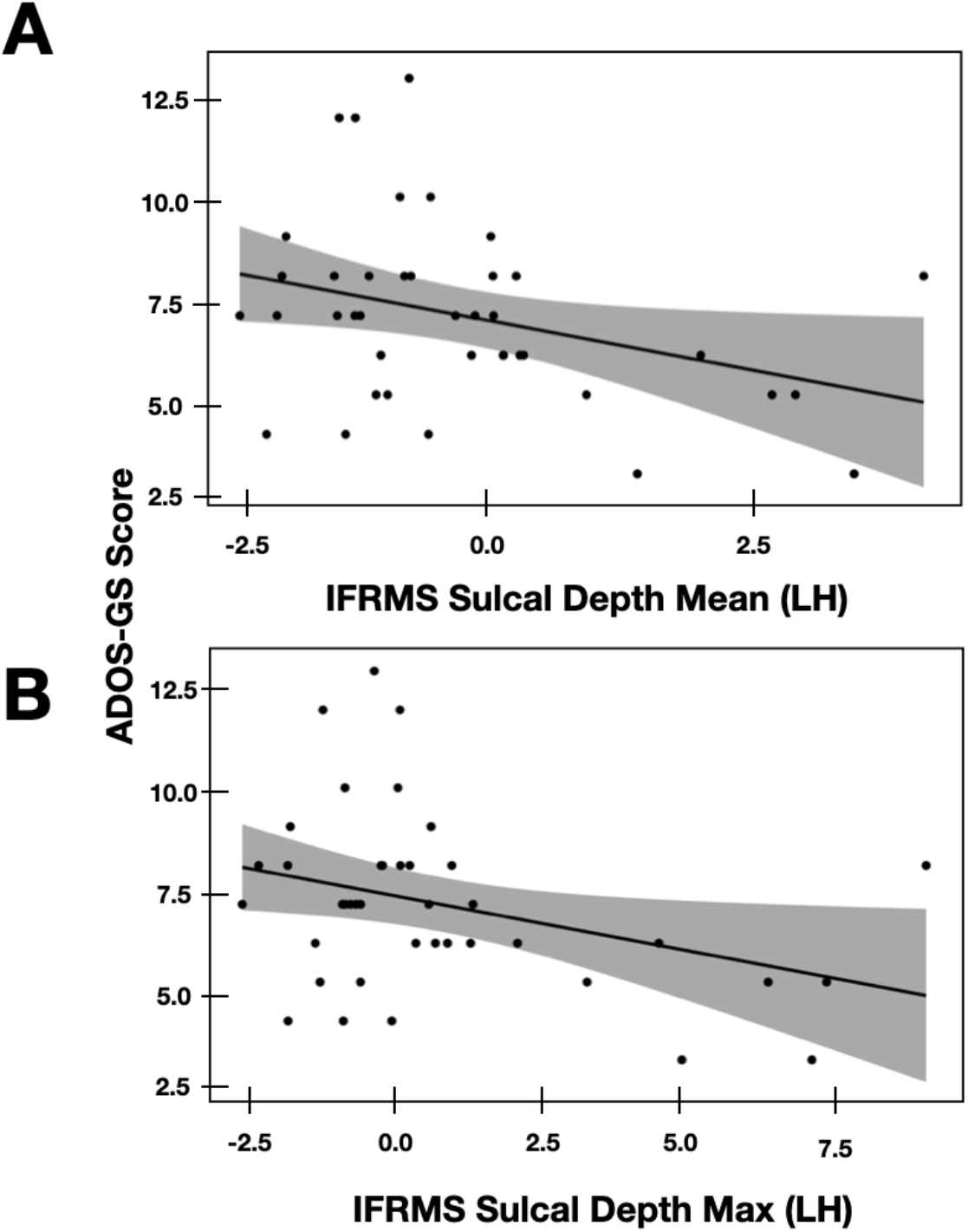
Sulcal depth of the IFRMS in the left hemisphere is negatively correlated with ADOS-GS scores. **A**. The mean sulcal depth of the IFRMS is negatively correlated with the ADOS-GS score (*r* = -0.33, *p* = 0.036). **B**. The max sulcal depth of the IFRMS is negatively correlated with the ADOS-GS score (*r* = -0.34, *p* = 0.031).

## Discussion

To our knowledge, the present measurements were the first to characterize tertiary sulci in PMC in ASD participants, and the second to do so in VTC (Ammons et al., 2021). Our analyses revealed morphological differences of VTC and PMC sulci between groups with four main findings. First and second, the largest effect was that individual differences in cortical thickness across all VTC and PMC sulci were larger in individuals with ASD compared to NTs – replicating and extending previous findings (Ammons et al., 2021) to other sulci in VTC aside from the MFS, as well as to PMC sulci. Third, the MFS was shorter in individuals with ASD compared to NTs, which is consistent with recent findings showing that the MFS is also shorter in individuals with developmental prosopagnosia compared to NTs (Parker et al., 2022). Fourth, correlation analyses between tertiary sulci in both regions revealed that some, but not all, tertiary sulcal morphological features of VTC and PMC are correlated in ASD individuals. Finally, we found that individual differences in IFRMS depth predicted individual differences in ADOS-GS scores. Below, we discuss these findings in the context of i) the relationship between a potential role of tertiary sulcal morphology and ASD, as well as other neurodevelopmental disorders, ii) underlying white matter architecture contributing to the relationships identified here such as potential white matter tracts connecting tertiary sulci in VTC and PMC, and iii) the limitations and future directions of the present study.

### On the role of tertiary sulci in neurodevelopmental disorders

Our findings build on a growing number of studies documenting the role of tertiary sulci in neurodevelopmental disorders (Ammons et al., 2021; Brun et al., 2016; Garrison et al., 2015; Garrison, Fernyhough, McCarthy-Jones, Simons, & Sommer, 2019; Hettwer et al., 2022; Parker et al., 2022; Powers, van Dyck, Garrison, & Corlett, 2020; Rollins et al., 2020). For example, in VTC, three recent studies found a relationship between anatomical features of either the fusiform gyrus at the location of the MFS, or the MFS morphology in particular, in different neurodevelopmental disorders. Specifically, Ammons et al. (2021) found morphological differences in the MFS of ASD participants when compared to NTs. In their analyses, they found that MFS cortical thickness was significantly correlated with the ability to interpret emotion and to infer mental states from facial features. Additionally, recent work examining morphological features of the MFS in developmental prosopagnosia revealed that the MFS was significantly shorter in developmental prosopagnosics compared to NTs, and that MFS length also predicted face perception ability (Parker et al., 2022). Finally, recent work (Hettwer et al., 2022) identified that the morphology of the right mid-fusiform gyrus was positively correlated with schizophrenia polygenic resilience scores overlapping with the location of the MFS at the group level. Thus, the combination of these findings indicates that MFS length is likely a transdiagnostic factor in both ASD and DP, and that future work performing morphological analyses at the level of individual participants in additional populations (such as individuals with schizophrenia) may further empirically support, as well as clarify, the role of MFS morphology in a growing number of neurodevelopmental disorders.

Beyond VTC, the present findings also incorporate novel PMC sulci for the first time and show that, like VTC (Ammons et al., 2021), individual differences in sulcal thickness across sulcal types are greater in individuals with ASD compared to NTs. Further, these findings revealed that some, but not all, morphological features of tertiary PMC and VTC sulci are correlated with one another in ASD individuals, suggesting that these correlated features of PMC and VTC may developmentally emerge at similar time points —a hypothesis which can be explored further in future studies.

Critically, the present findings also show a relationship between the morphology of a newly identified tertiary sulcus in PMC (the IFRMS) and behavior for the first time. Specifically, we show that individual differences in IFRMS depth predict individual differences in ADOS score in which the deeper the sulcus, the lower the ADOS score. The fact that only sulcal depth, but not other morphological features, were correlated with ADOS score is consistent with recent results showing a relationship between sulcal depth in lateral prefrontal cortex and higher-level aspects of cognition such as relational reasoning (Voorhies et al., 2021) and working memory (Yao, Voorhies, Miller, Bunge, & Weiner, 2022) in a developmental cohort, as well as inferior frontal cortex in ASD individuals. In terms of the latter, Brun et al. (2016) found that sulcal pit depth in the left hemisphere of the ascending ramus of the lateral fissure was correlated with social communication behavior in ASD individuals.

Complementing the present findings that targeted face processing, Ammons and colleagues (2021) focused on tasks related to theory of mind [Autism Spectrum Quotient (AQ) and the Reading the Mind in the Eyes task (RMIE)] in order to examine the relationship between sulcal morphology and social behaviors. We utilized ADOS-GS scores instead since this test presents participants with tasks explicitly designed to quantify behavioral components of facial processing such as facial expressions and eye orientation during social interactions. Ammons and colleagues (2021) found that the morphology of the left MFS was correlated with performance on the RMIE. Although the morphology of the MFS was not related to ADOS-GS, our findings did extend the relationship between sulcal morphology and cognition in ASD to PMC sulci. Specifically, differences in the sulcal depth of the IFRMS were related to differences in ADOS-GS scores. Thus, building on the work from Ammons and colleagues (2021) and Brun and colleagues (2016), we further show that this relationship between left hemisphere sulcal morphology and behavior in ASD extends to PMC.

Associations between individual differences in sulcal morphology and individual differences in behavior is not limited to visual processing or ASD. For example, in the medial prefrontal cortex, Garrison et al. (2015) revealed that there was a reduction in the length of the paracingulate sulcus in patients with schizophrenia who hallucinated compared to non-hallucinating patients and NTs, which has been replicated (Garrison et al., 2019; Powers et al., 2020; Rollins et al., 2020). Taken together, these findings emphasize that tertiary sulcal morphology is correlated with behavior in multiple regions in multiple neurodevelopmental disorders.

### White matter tracts connecting VTC and PMC tertiary sulci?

Considering the morphological relationship between the IFRMS in PCC and the MFS in VTC in ASD participants but not NTs (Figure 6), as well as theories linking sulcal depth to underlying white matter (Van Essen, 2020; Van Essen, 1997; Welker, 1990), we hypothesize that presently unidentified corticocortical projections between the MFS and the IFRMS may be shorter in ASD participants than NTs. These ideas are in line with recent studies showing that a vertical component of the cingulum bundle connects PMC with VTC. For instance, Jones et al. (2013) found that a minority of cingulum bundle fibers were divided into three sections based on the trajectory of their fibers, one of which connects the cingulate with ventral temporal regions. Specifically, Jones and colleagues (2013) referred to this bundle as the “restricted parahippocampal cingulum”, and recognized that further examination was required to better understand its functional value as well as to better characterize the specific projections of the bundle. Shortly thereafter, Wu et al (2016) dissected the cingulum bundle into five segments, one of which they referred to as Cingulum Bundle V (CB-V), which also connected the PMC to VTC. Similar observations were also made by Skandalakis et al (2020), who noted that the functional role, as well as clinical implications of this bundle, were presently lacking. Thus, future studies can test the following hypotheses, if: i) CB-V connects the IFRMS to the MFS and ii) features of CB-V are different in ASD and NT participants.

### Future directions and limitations

The findings from the present study lead to at least three additional studies moving forward. First, while the present study i) compares sulcal morphology between ASD participants and NTs and ii) examines the relationship between sulcal morphology and cognition in ASD participants, it did not also consider functional representations. For example, the IFRMS has recently been shown to identify a region of the cognitive control network (Willbrand et al., in press) and face-selective regions have been recently identified in PMC (Silson, Steel, Kidder, Gilmore, & Baker, 2019; Woolnough et al., 2020), while the MFS identifies functional transitions in many functional maps and also, predicts the location of face-selective regions (Grill-Spector & Weiner, 2014; Weiner, 2019). Thus, future studies can also quantify the relationship of sulcal morphology to these different functional representations in ASD participants and NTs, and then compare this relationship between groups. Second, as mentioned at the end of the previous section, an analysis of integrity differences in the CB-V between ASD participants and NTs using DWI would provide insights regarding clinical implications of this bundle. Third, the difference between hemispheres noted in a number of our findings identifies a need for future studies to explore lateralization differences between ASD participants and NTs in both the anatomical, as well as the functional, organization of sulcal morphology in VTC and PMC. In terms of limitations, a main limiting factor of our behavioral analyses is that the ABIDE dataset did not provide ADOS-GS scores for NTs. Therefore, the results of our analyses considering behavior were limited to individual differences in performance among those with ASD, but not NTs.

### Conclusion

Building on previous work showing that VTC sulcal morphology differs between ASD participants and NTs (Ammons et al., 2021), our findings show that PMC sulcal morphology also differs between ASD and NT participants. Some morphological metrics were also correlated with performance on a facial processing task (ADOS-GS), revealing that sulcal depth of the IFRMS was correlated with individual differences in cognition. These findings are consistent with a growing body of work showing that individual differences in the morphology of tertiary sulci are related to individual differences in behavior. Our findings are encouraging for the study of tertiary sulci in ASD in other cortical expanses and other neurodevelopmental disorders, which can be explored in future studies.

## Acknowledgments

This research was supported by NSF CAREER Award 2042251 (Weiner). Funding for original data collection and curation for the New York University sample was provided by NIH (K23MH087770; R21MH084126; R01MH081218; R01HD065282), Autism Speaks, The Stavros Niarchos Foundation, The Leon Levy Foundation, and an endowment provided by Phyllis Green and Randolph Cō wen. Funding for original data collection and curation for the Georgetown University sample was provided by NIMH MH084961, Intellectual and Developmental Disabilities Research Center, and Children’s National Medical Center HD040677-07. We thank Linda Wilbrecht for providing feedback on the analysis and manuscript in its initial stages.

