## Supplementary tables and figures for "Ventral temporal and posteromedial sulcal morphology in autism spectrum disorder"

| | NT (*n* = 50) | ASD (*n* = 50) | *T* value | *P* value |
|---|---|---|---|---|
| **Gender** | 39 M, 11 F | 44 M, 6 F | - | - |
| **Age (years)** | 10.23 (± 2.89; 5.89-23.81) | 10.16 (± 5.75; 5.32-34.76) | -0.162 | 0.871 |
| **VIQ** | 119.63 (± 14.82; 85-143) | 101.94 (± 16.79; 68-135) | -5.53 | <.001** |
| **PIQ** | 101.94 (± 16.79; 68-135) | 102.19 (± 19.08; 67-147) | -3.503 | <.001** |
| **FIQ** | 118.9 (± 14.56; 91-144) | 102.73 (± 18.58; 67-138) | -4.794 | <.001** |

**Supplementary Table 1. Participant Demographics*.*** *Note.* (± Standard deviation; range), FIQ: Full Intelligence Quotient; PIQ: Performance IQ; VIQ: Verbal IQ. **P < 0.001.

| Sulcus | Region | ASD LH # | ASD LH % | ASD RH # | ASD LH % | NT LH # | NT LH % | NT RH # | NT RH % |
|---|---|---|---|---|---|---|---|---|---|
| MFS | VTC | 50 | 1 | 50 | 1 | 50 | 1 | 50 | 1 |
| CoS | VTC | 50 | 1 | 50 | 1 | 50 | 1 | 50 | 1 |
| OTS | VTC | 50 | 1 | 50 | 1 | 50 | 1 | 50 | 1 |
| MCGS | PMC | 50 | 1 | 50 | 1 | 50 | 1 | 50 | 1 |
| POS | PMC | 50 | 1 | 50 | 1 | 50 | 1 | 50 | 1 |
| SPLS | PMC | 50 | 1 | 50 | 1 | 50 | 1 | 50 | 1 |
| PRCULS | PMC | 50 | 1 | 50 | 1 | 50 | 1 | 50 | 1 |
| PRCUS-p | PMC | 50 | 1 | 50 | 1 | 50 | 1 | 49 | 0.98 |
| PRCUS-i | PMC | 50 | 1 | 50 | 1 | 50 | 1 | 50 | 1 |
| PRCUS-a | PMC | 50 | 1 | 50 | 1 | 50 | 1 | 49 | 0.98 |
| SPS | PMC | 46 | 0.92 | 46 | 0.92 | 42 | 0.84 | 42 | 0.84 |
| IFRMS | PMC | 50 | 1 | 50 | 1 | 49 | 0.98 | 49 | 0.98 |
| SSPLS | PMC | 29 | 0.58 | 26 | 0.52 | 19 | 0.38 | 26 | 0.52 |
| ICGS-p | PMC | 24 | 0.48 | 29 | 0.58 | 21 | 0.42 | 33 | 0.66 |

**Supplementary Table 2. Sulcal Incidence Rates.** *Note.* '#' pertains to the number of sulci per group, in which the total possible is 50, percentage (%) is out of 1.

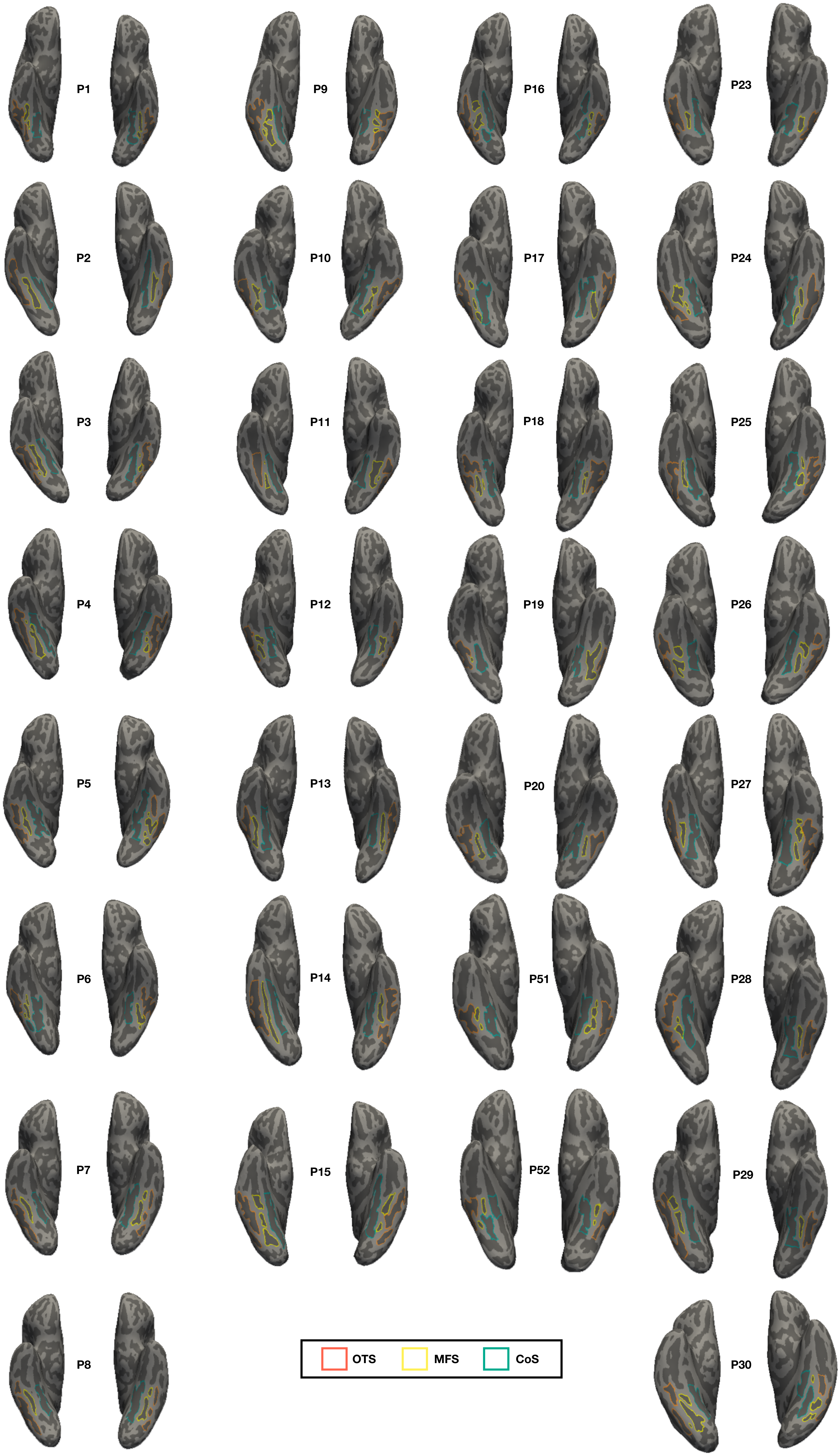
P1
P9
P16
P23
P2
P10
P17
P24
P3
P11
P18
P25
P4
P12
P19
P26
P5
P13
P20
P27
P6
P14
P51
P28
P7
P15
P52
P29
P8
OTS
MFS
CoS
P30

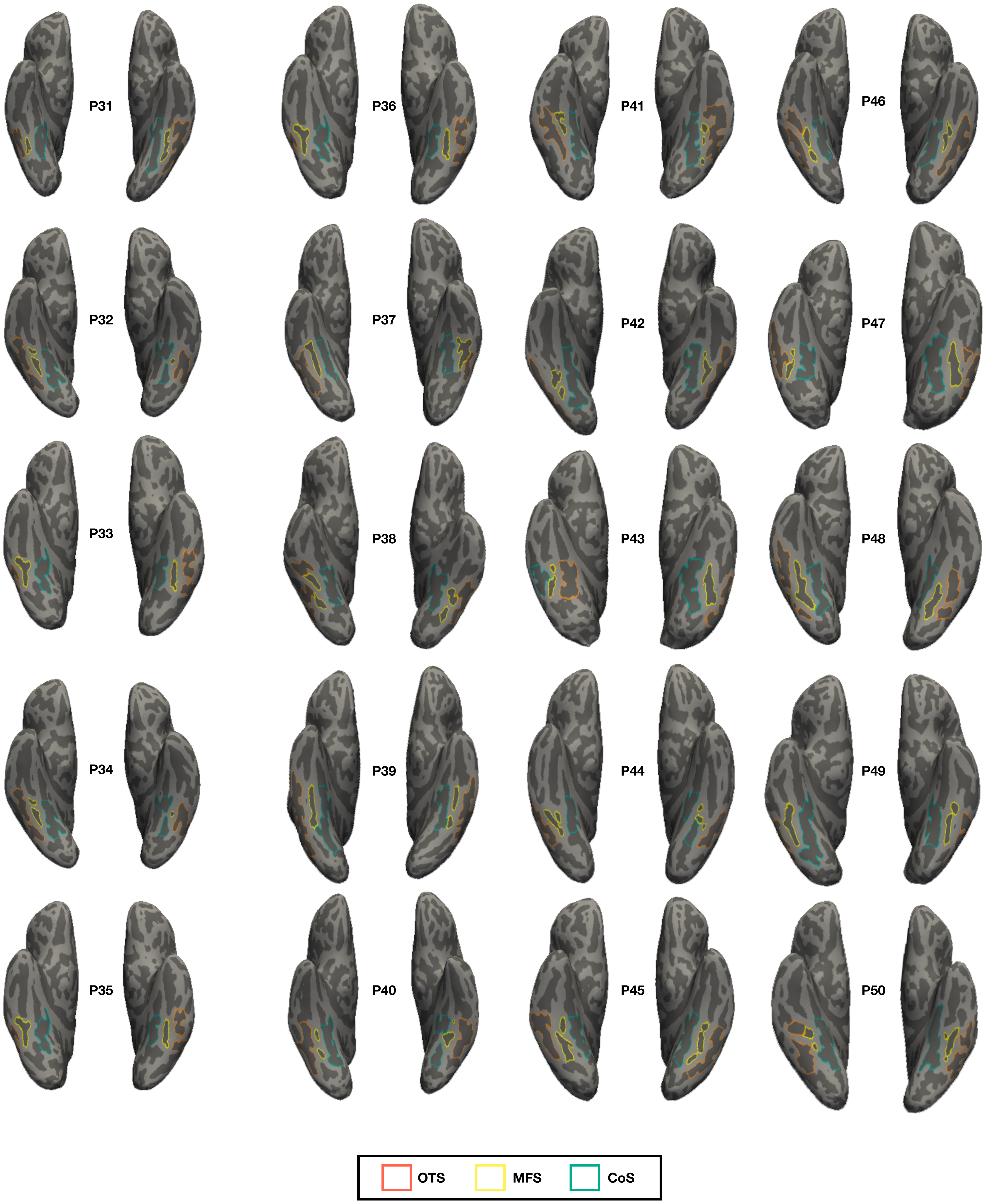


**Supplementary Figure 1. Manual VTC sulcal labels in the left and right hemispheres of each participant in the NT sample.** Each sulcus is displayed on the inflated cortical surface in FreeSurfer 6.0.0 and is colored according to the key at the top. Each hemisphere contains three sulci: the mid-fusiform sulcus (MFS), occipitotemporal sulcus (OTS), and collateral sulcus (CoS).

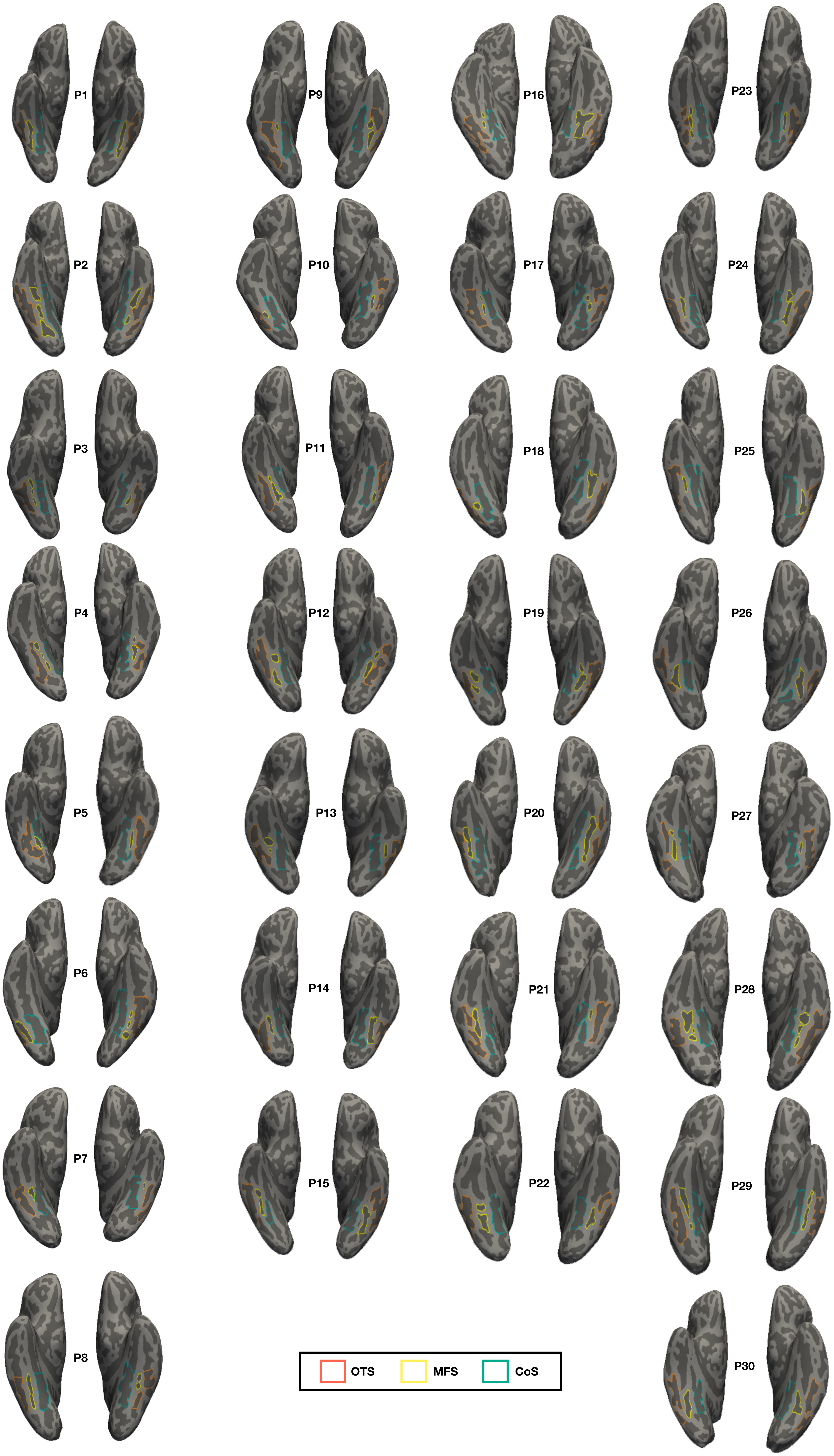
P1
P9
P16
P23
P2
P10
P17
P24
P3
P11
P18
P25
P4
P12
P19
P26
P5
P13
P20
P27
P6
P14
P21
P28
P7
P15
P22
P29
P8
P30
OTS
MFS
CoS

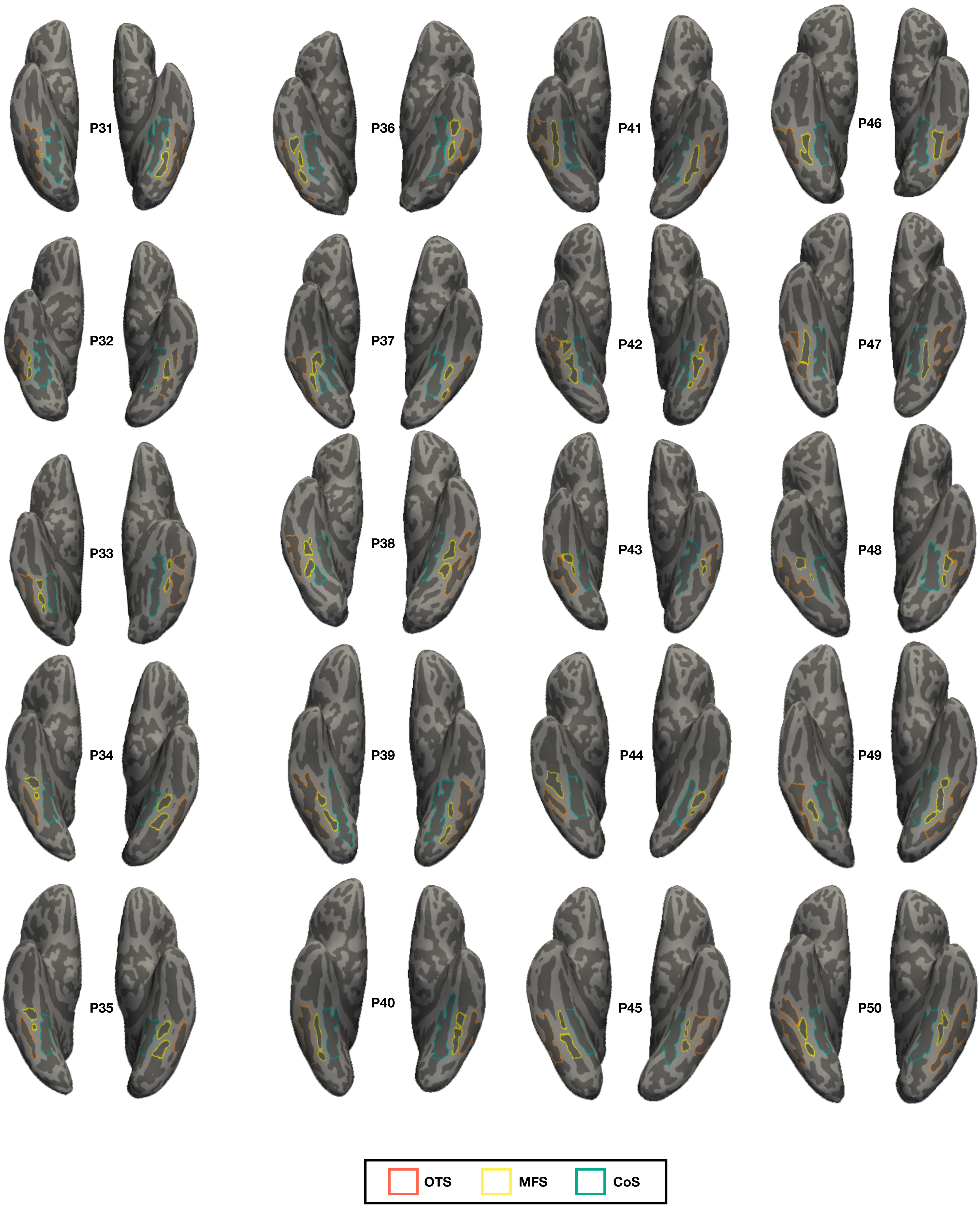


**Supplementary Figure 2. Manual VTC sulcal labels in the left and right hemispheres of each participant in the ASD sample.** Each sulcus is displayed on the inflated cortical surface in FreeSurfer 6.0.0 and is colored according to the key at the top. Each hemisphere contains three sulci: the mid-fusiform sulcus (MFS), occipitotemporal sulcus (OTS), and collateral sulcus (CoS).

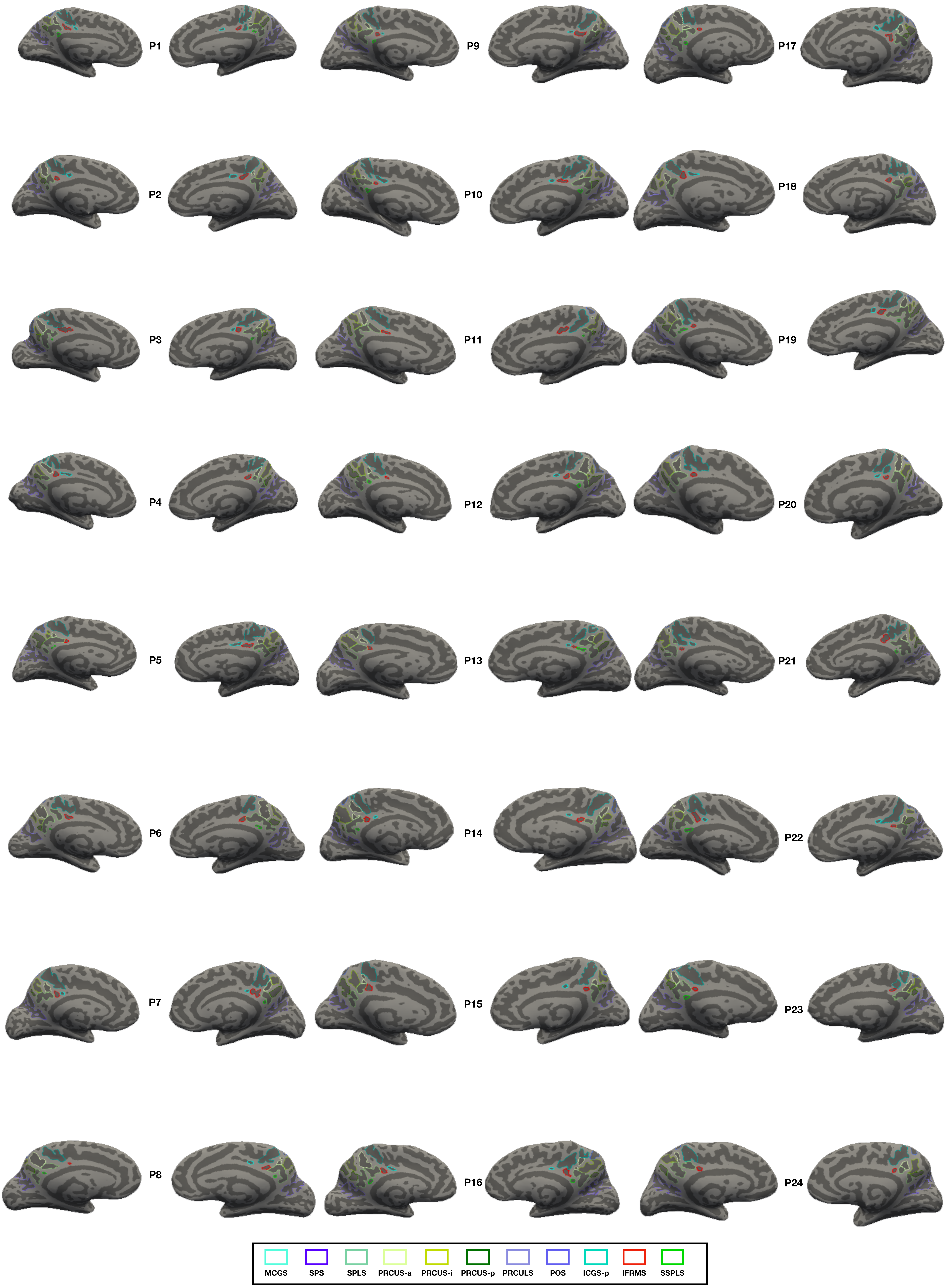

P1
P2
P3
P4
P5
P6
P7
P8
P9
P10
P11
P12
P13
P14
P15
P16
P17
P18
P19
P20
P21
P22
P23
P24
MCGS
SPS
SPLS
PRCUS-a
PRCUS-i
PRCUS-p
PRCULS
POS
ICGS-p
IFRMS
SSPLS

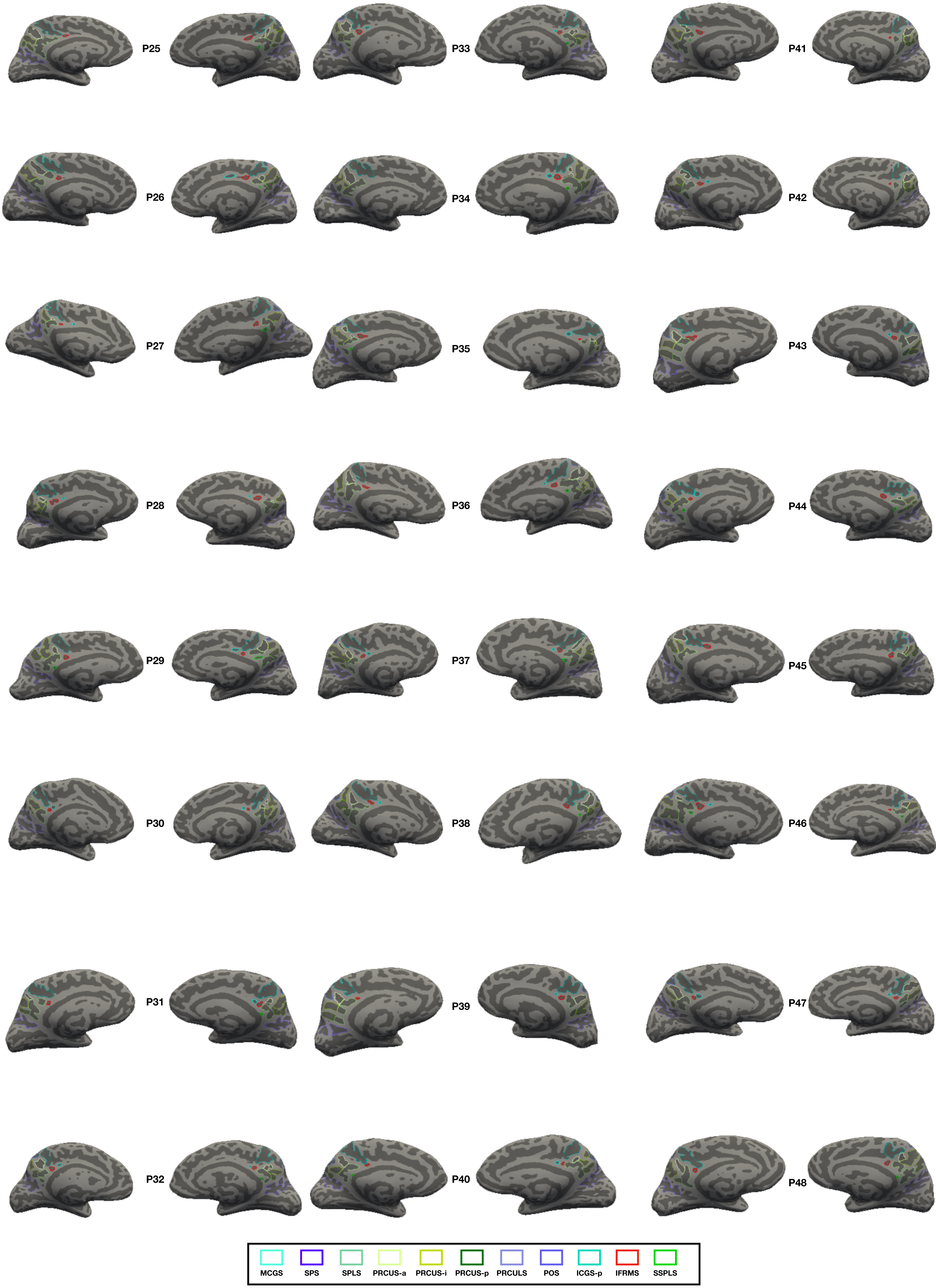

P25
P33
P41
P26
P34
P42
P27
P35
P43
P28
P36
P44
P29
P37
P45
P30
P38
P46
P31
P39
P47
P32
P40
P48
MCGS
SPS
SPLS
PRCUS-a
PRCUS-i
PRCUS-p
PRCULS
POS
ICGS-p
IFRMS
SSPLS

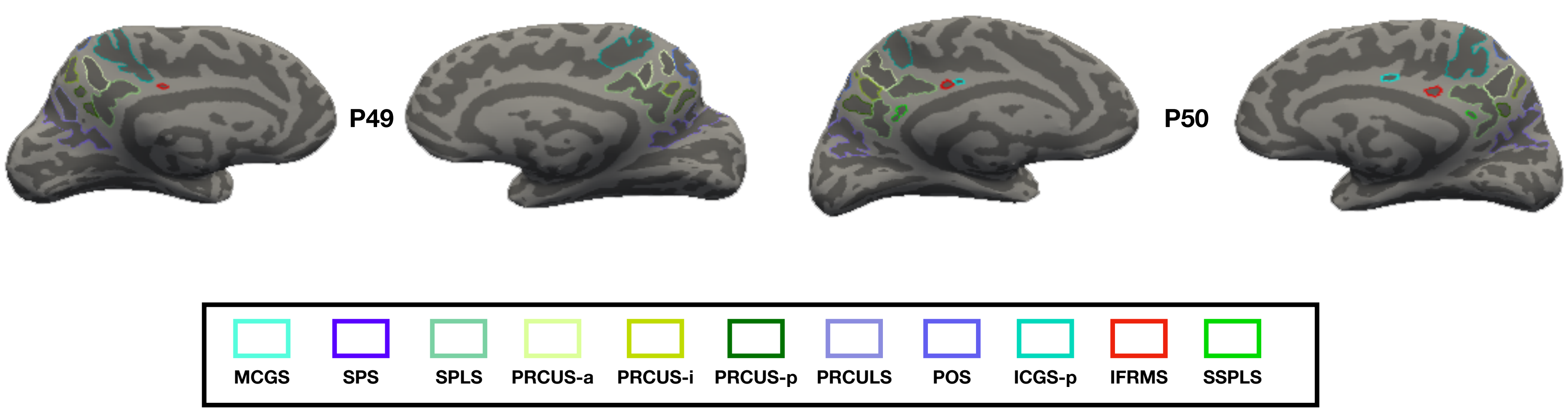


**Supplementary Figure 3. Manual PMC sulcal labels in the left and right hemispheres of each participant in the NT sample.** Each sulcus is displayed on the inflated cortical surface in FreeSurfer 6.0.0 and is colored according to the key at the top. Each hemisphere contains at least eight sulci (from posterior to anterior): pos, prculs, prcus-p, prcus-i, prcus-a, spls, mcgs, and ifrms. The sps, sspls, and icgs-p are all variably present.

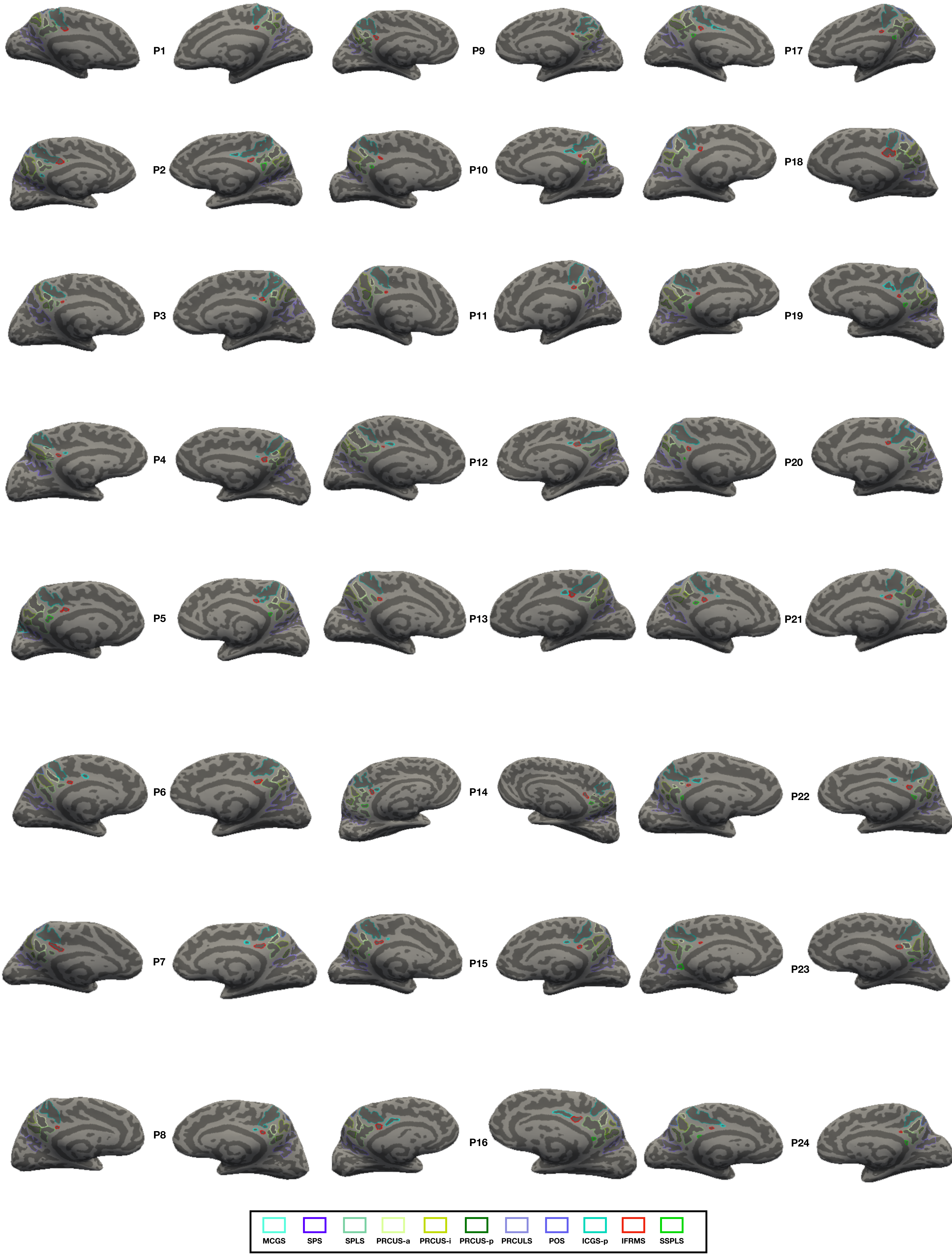
P1
P9
P17
P2
P10
P18
P3
P11
P19
P4
P12
P20
P5
P13
P21
P6
P14
P22
P7
P15
P23
P8
P16
P24
MCGS
SPS
SPLS
PRCUS-a
PRCUS-i
PRCUS-p
PRCULS
POS
ICGS-p
IFRMS
SSPLS

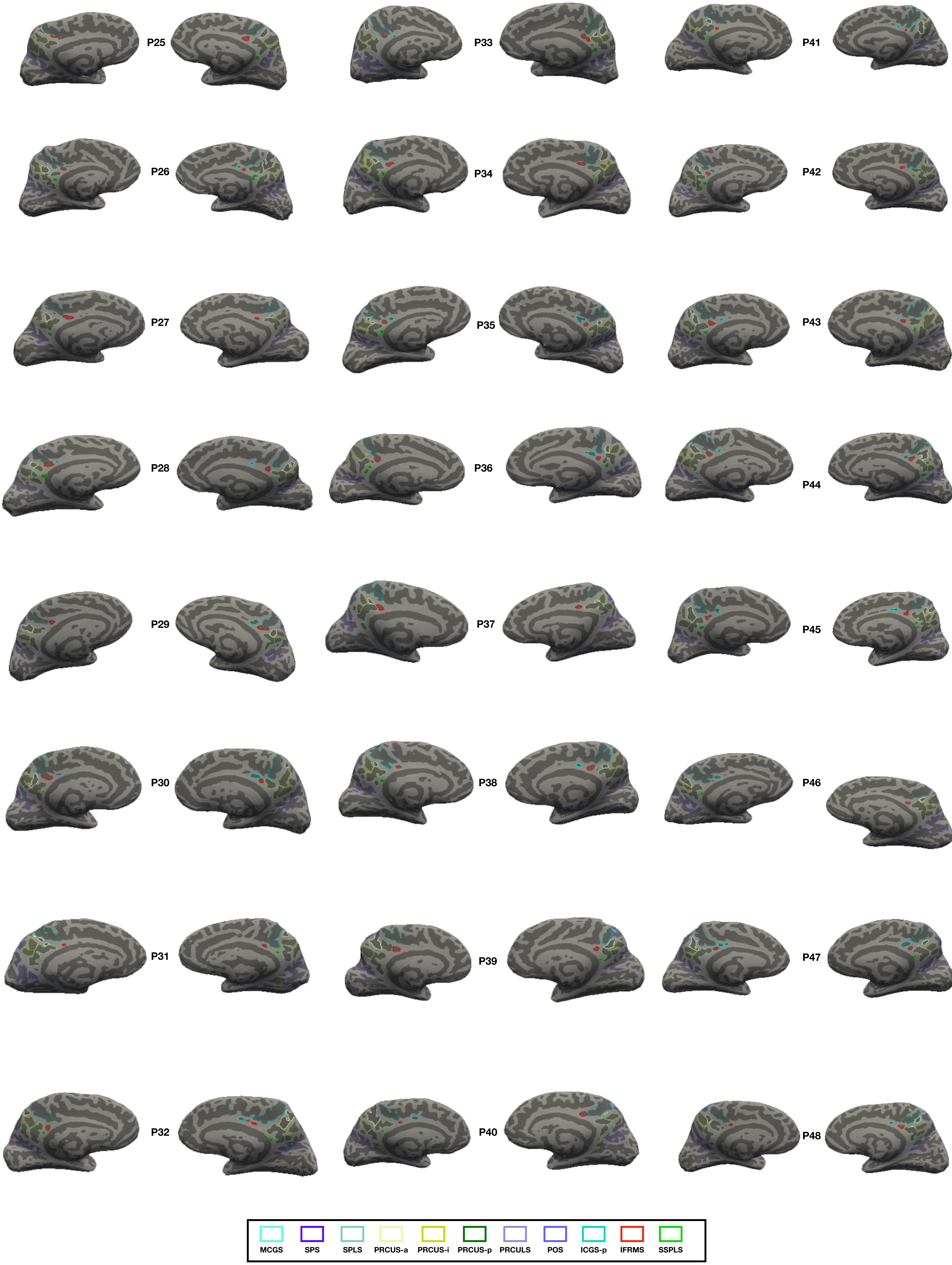

P25
P33
P41
P26
P34
P42
P27
P35
P43
P28
P36
P44
P29
P37
P45
P30
P38
P46
P31
P39
P47
P32
P40
P48
MCGS
SPS
SPLS
PRCUS-a
PRCUS-i
PRCUS-p
PRCULS
POS
ICGS-p
IFRMS
SSPLS

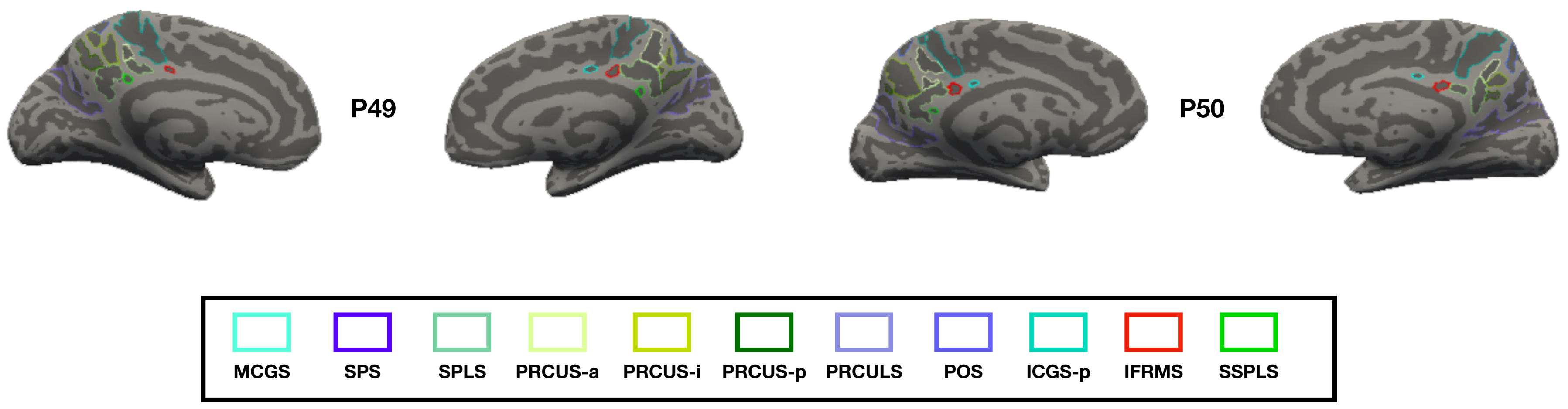


**Supplementary Figure 4. Manual PMC sulcal labels in the left and right hemispheres of each participant in the ASD sample.** Each sulcus is displayed on the inflated cortical surface in FreeSurfer 6.0.0 and is colored according to the key at the top. Each hemisphere contains at least eight sulci (from posterior to anterior): pos, prculs, prcus-p, prcus-i, prcus-a, spls, mcgs, and ifrms. The sps, sspls, and icgs-p are all variably present.

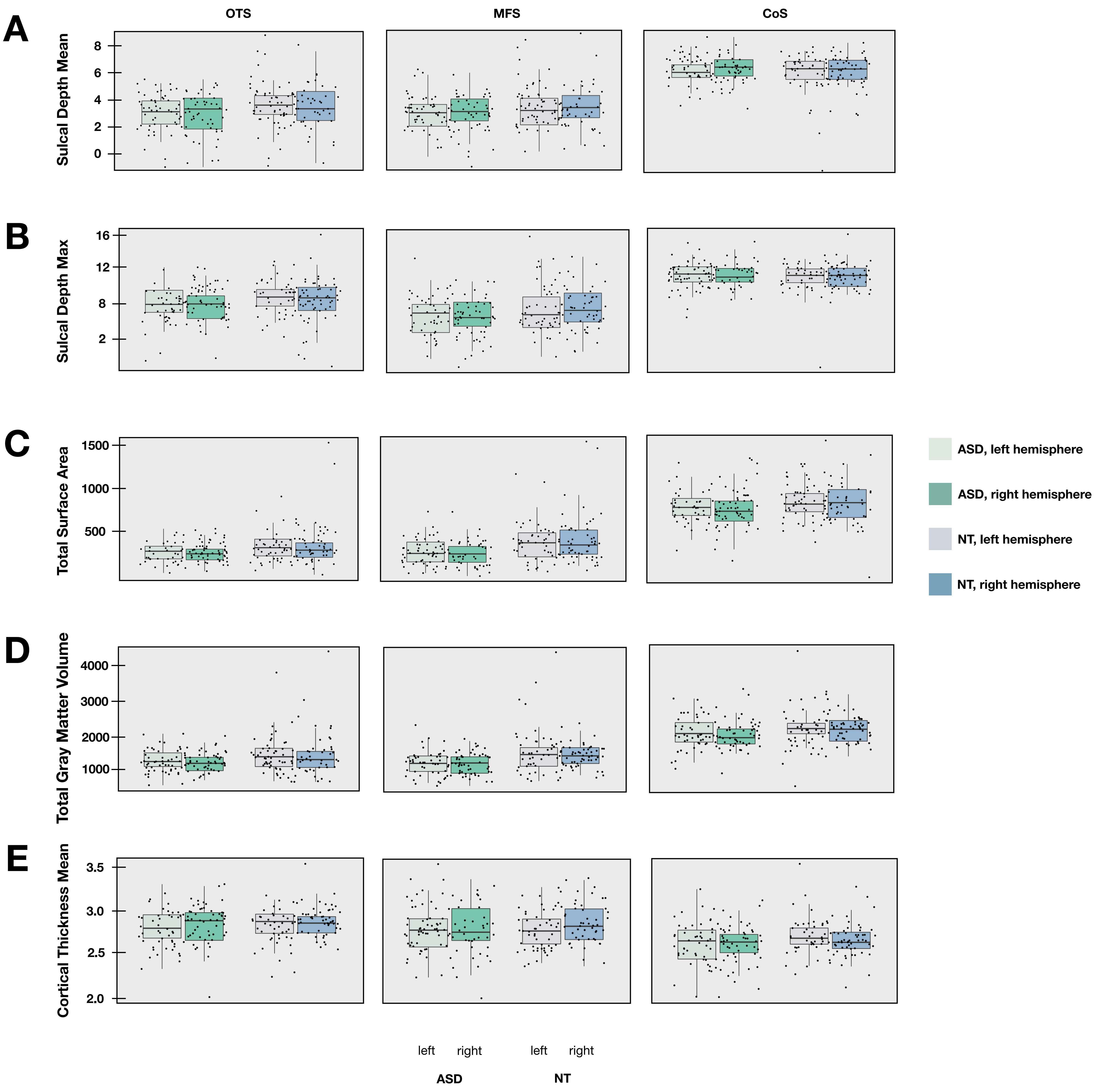


**Supplementary Figure 5. Non-significant features across VTC sulci.** The above box plots indicate all non-significant features (±quartile) of all VTC sulci. The left two plots indicate the left and right hemispheres of the ASD group and the right two plots indicate the left and right hemispheres of the NT group, respectively. **A.** Sulcal depth mean **B.** Sulcal depth max **C.** Total surface area **D.** Gray matter volume **E.** Cortical thickness mean.

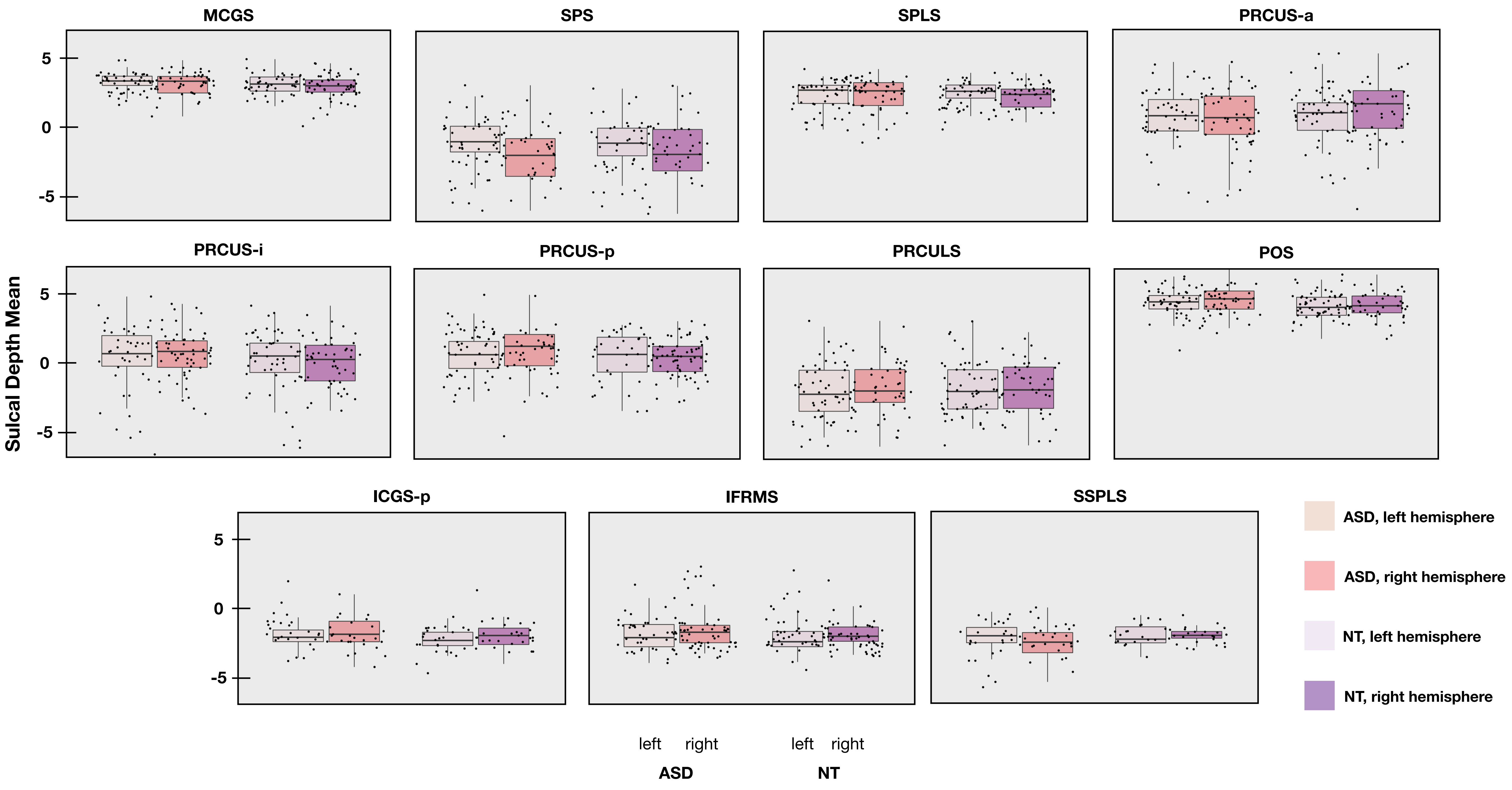


**Supplementary Figure 6. Non-significant features across PMC sulci: Mean sulcal depth.** The above box plots indicate sulcal depth mean (±quartile) of PMC sulci. The left two plots indicate the left and right hemispheres of the ASD group and the right two plots indicate the left and right hemispheres of the NT group, respectively.

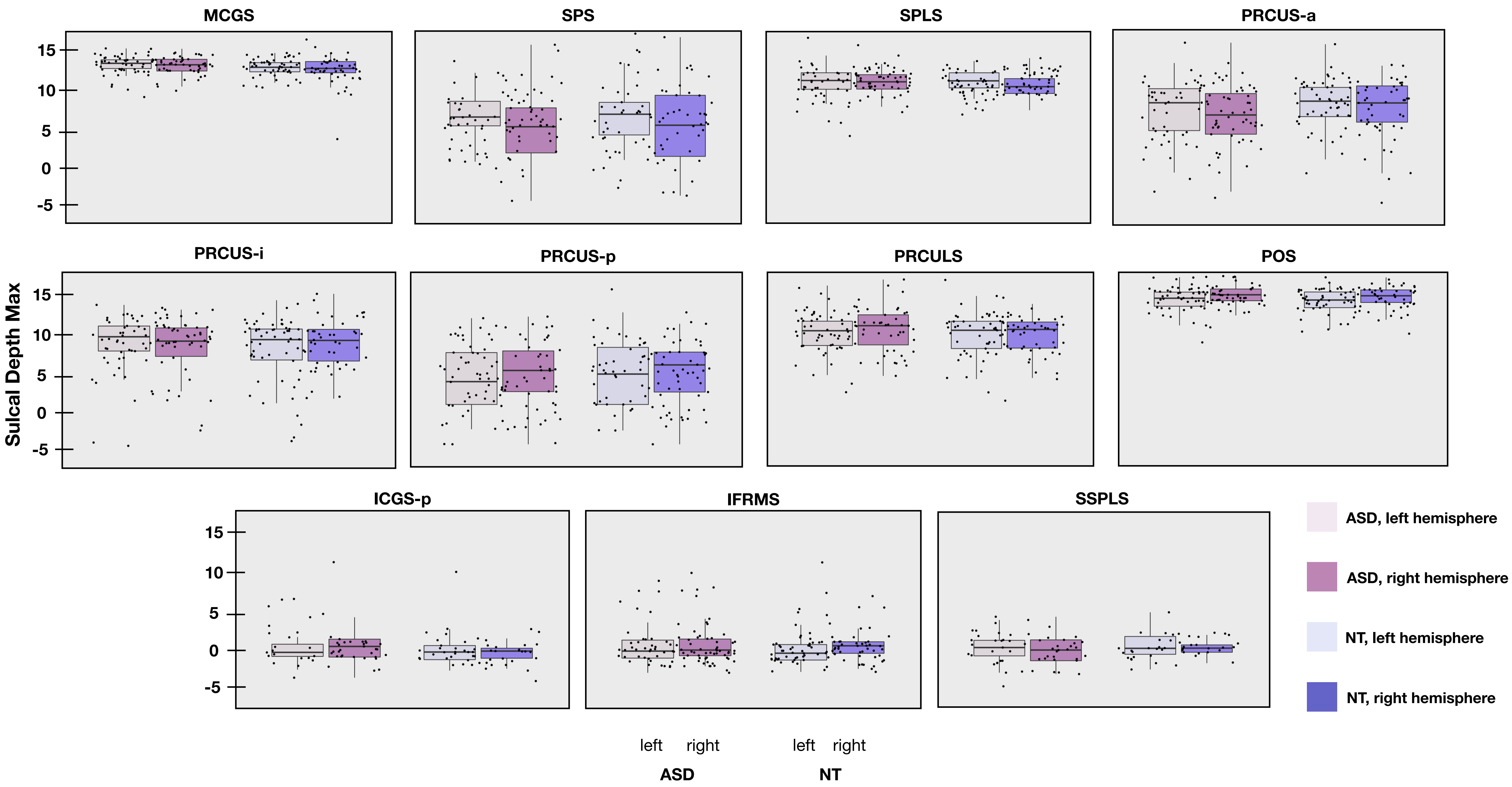


**Supplementary Figure 7. Non-significant features across PMC sulci: Max sulcal depth.** The above box plots indicate the sulcal depth max (±quartile) of all PMC sulci. The left two plots indicate the left and right hemispheres of the ASD group and the right two plots indicate the left and right hemispheres of the NT group, respectively.

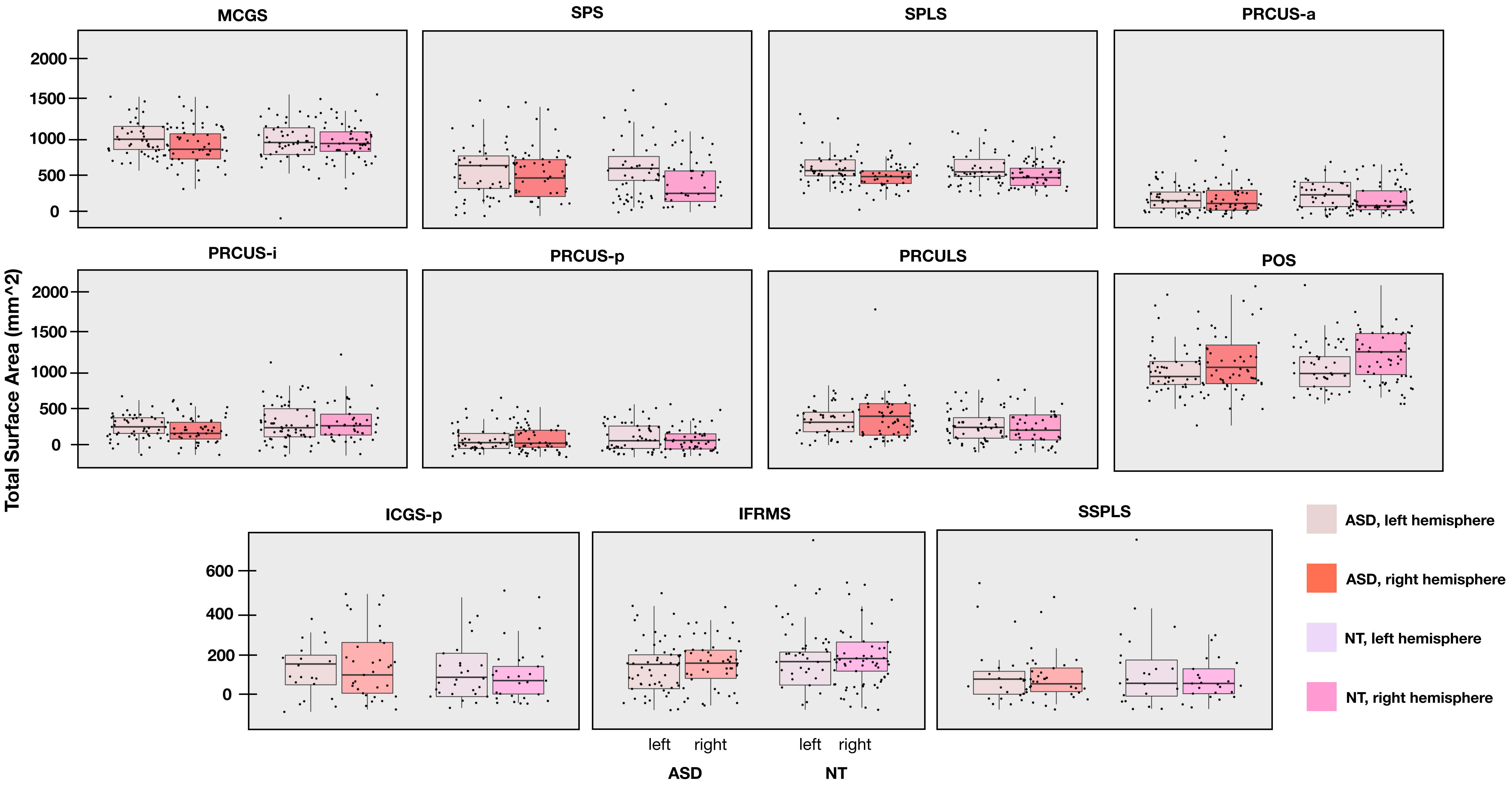


**Supplementary Figure 8. Non-significant features across PMC sulci: Surface area.** The above box plots indicate the total surface area (±quartile) of all PMC sulci. The left two plots indicate the left and right hemispheres of the ASD group and the right two plots indicate the left and right hemispheres of the NT group, respectively.

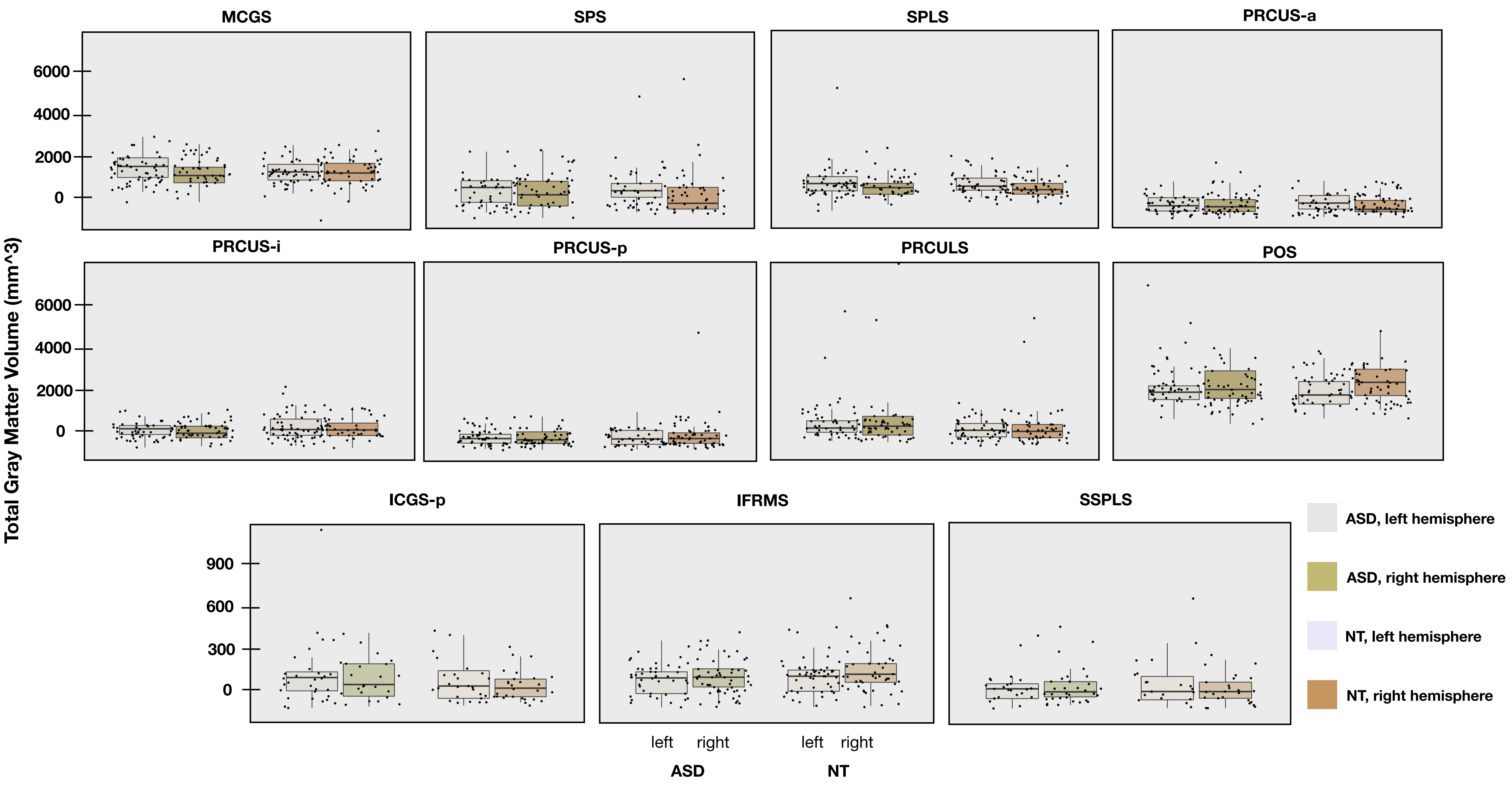


**Supplementary Figure 9. Non-significant features across PMC sulci: Gray matter volume.** The above box plots indicate the total gray matter volume (±quartile) of all PMC sulci. The left two plots indicate the left and right hemispheres of the ASD group and the right two plots indicate the left and right hemispheres of the NT group, respectively.

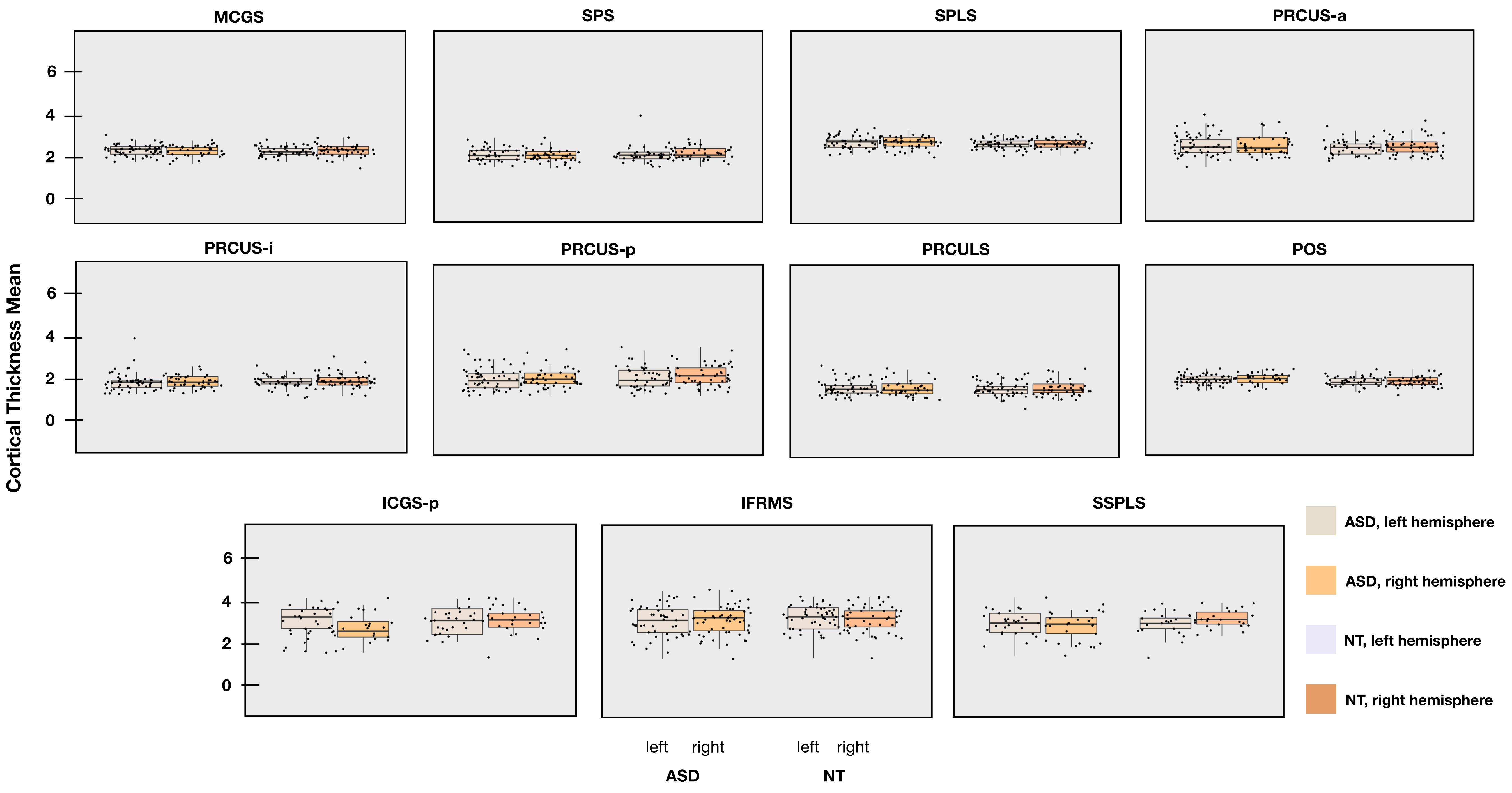


**Supplementary Figure 10. Non-significant features across PMC sulci: Cortical thickness.** The above box plots indicate the cortical thickness mean (±quartile) of all PMC sulci. The left two plots indicate the left and right hemispheres of the ASD group and the right two plots indicate the left and right hemispheres of the NT group, respectively.
